# Jumbo circular extrachromosomal elements of methane-oxidizing archaea with variably extensive metabolic and defense gene repertoires

**DOI:** 10.64898/2026.01.21.700959

**Authors:** Ling-Dong Shi, Bethany C. Kolody, Shuai Wang, Luis E. Valentin-Alvarado, Shufei Lei, Rohan Sachdeva, Jillian F. Banfield

**Affiliations:** Innovative Genomics Institute, University of California, Berkeley, CA, USA; Earth and Planetary Science, University of California, Berkeley, CA, USA; Biomedicine Discovery Institute, Monash University, Melbourne, Victoria, Australia; Environmental Science, Policy and Management, University of California, Berkeley, CA, USA

## Abstract

Archaeal extrachromosomal elements (ECEs) are arguably the least well understood of all genetic elements, and few have > 200 kbp (“jumbo”) genomes. Here, we report circular, jumbo ECEs with genomes of up to 535 kbp in length that associate with anaerobic methane-oxidizing *Methanoperedens* archaea. Notably, a 409-kbp genome related to jumbo ECEs is integrated into a subset of the ∼4.2 Mbp *Methanoperedens* chromosomes at the tRNA-Asp genes. This represents the largest integrative element in Archaea and supports the jumbo ECE - host association. Multiple genome alignment and phylogenetic analyses suggest that the large sizes were developed by extensive DNA acquisition from *Methanoperedens*. The newly identified ECEs encode, and in some cases express, metabolic genes such as tetrahydromethanopterin S-methyltransferase exclusively involved in methane oxidation, and genes for nitrogen and sulfur compound transformations. Also encoded are defense systems, some of which are absent in hosts, such as hybrid Type I/Type III-A CRISPR-Cas systems. In contrast to viruses and plasmids, they have host-like replication machinery and occur at stable copy ratios of 1.44 ± 0.24 : 1 to the host. Overall, our results reveal a spectrum of jumbo ECEs of *Methanoperedens*, ranging from plasmid-like to minichromosome-like.

## Introduction

All bacteria and archaea likely harbor extrachromosomal genetic elements (ECEs), including viruses, plasmids, and unclassified elements^1–4^. Some ECEs can integrate into host chromosomes (iMGEs) and subsequently excise^5–8^. Others, exemplified by plasmids, establish long-term coexistence within the host cell. ECEs and iMGEs profoundly shape host physiology, ecological fitness, and evolutionary trajectories. Viruses can reprogram host intracellular processes and induce host morphotypic changes^9–11^. Plasmids can confer advantageous traits to their hosts, such as genes that enable hosts to respond to environmental stresses (e.g., antibiotics)^12–14^. Extensive interactions between ECEs and hosts, during long-term coexistence, may drive their coevolution^15,16^.

Most prokaryotic ECEs with genomes of > 200 kbp in length (“jumbo”) have been found in Bacteria^17–19^. Jumbo archaeal ECEs include circular minichromosomes of Haloarchaea, sometimes designated megaplasmids^20–23^, one undescribed ∼285 kbp putative plasmid of *Methanomethylovorans hollandica* (CP003363), and Borgs with linear genomes up to 1.1 Mb in length that are predicted to replicate in uncultured, anaerobic methane-oxidizing *Methanoperedens* archaea^24^.

Most archaeal ECEs, particularly those with small genomes, encode genes primarily related to DNA processing and few involved in metabolic pathways^25–28^. The exception is Borgs, which encode diverse genetic inventories for energy generation and core metabolism, including genes that encode the methyl-coenzyme M reductase (MCR) complex, genes for the TCA cycle, and genes for polyhydroxybutyrate production^24^. Here, we bring to light novel archaeal circular and integrated ECEs with 233 – 535 kbp genomes that encode numerous metabolic genes. Reconstruction and curation of complete ECE genomes enabled confident analyses of their genome architecture and genetic repertoires. The metabolic gene inventory and genome features suggest that these ECEs are on a continuum between plasmids and minichromosomes.

## Results

### Newly discovered jumbo ECEs are associated with *Methanoperedens*

To identify jumbo ECEs of archaea, we used a combination of long- and short-read sequencing to investigate wetland soil in California, a site that is known to host diverse groups of archaea, including methanogens, methanotrophs, Asgard archaea, and their associated extrachromosomal elements^8,24,28,29^. We sought putatively circular PacBio long-read derived genomes with lengths > 200 kbp, not classifiable as organisms (given the lack of rRNAs and ribosome components), and where proteins with identifiable matches were most similar to those of Archaea (see Methods). Three seemingly circular genomes were recovered, and their circularization was confidently verified by PacBio reads (Supplementary Fig. 1). Illumina reads were used to ensure base call support of the PacBio-derived genomes at every position. These circular complete genomes were then used to identify three related but partial sequences that were further curated to completion using PacBio long reads and Illumina short reads. All six genomes are circular and > 200 kbp in length (Supplementary Table 1). In addition, we identified a 409-kbp related element from rice field soil that is integrated into a *Methanoperedens* genome (Mp_4-22MB_43_25) at a tRNA-Asp gene (Supplementary Table 2). The element is flanked by 14-bp direct repeats: one is inside the integrated ECE and the other is on the host chromosome (Supplementary Fig. 2). Within the ECE and adjacent to the integration site is a tyrosine-type integrase gene. The integration clearly establishes *Methanoperedens* archaea as the host of the 409-kbp element, and likely for the other six circular > 200 kbp ECEs. Seven related but smaller, complete circular genomes (53.3 - 157.3 kbp in length) were also recovered (Supplementary Table 1).

To test for linkages between *Methanoperedens* archaea and the ECEs, we compared the proteins of the 14 ECEs to those of organisms in public databases and found that ∼62% of the proteins had best matches to those of *Methanoperedens* (see Methods). The fraction of proteins with best matches to *Methanoperedens* increased with increasing length of ECE genomes (Fig. 1). We also identified identical or near-identical transposons, comprising transposases and terminal inverted repeats, shared between 11 of the ECE genomes and *Methanoperedens* genomes (Supplementary Fig. 3). Another two ECEs (87kb_39_4 and 77kb_39_7) share 1,866-bp and 1,680-bp regions that include transposases (but with no identifiable terminal inverted repeats) and have ≤ 3 bp mismatches with regions in *Methanoperedens* genomes. This level of sequence similarity is strongly suggestive of recent DNA transfer and supports the association of ECEs with *Methanoperedens*.

**Figure 1.**
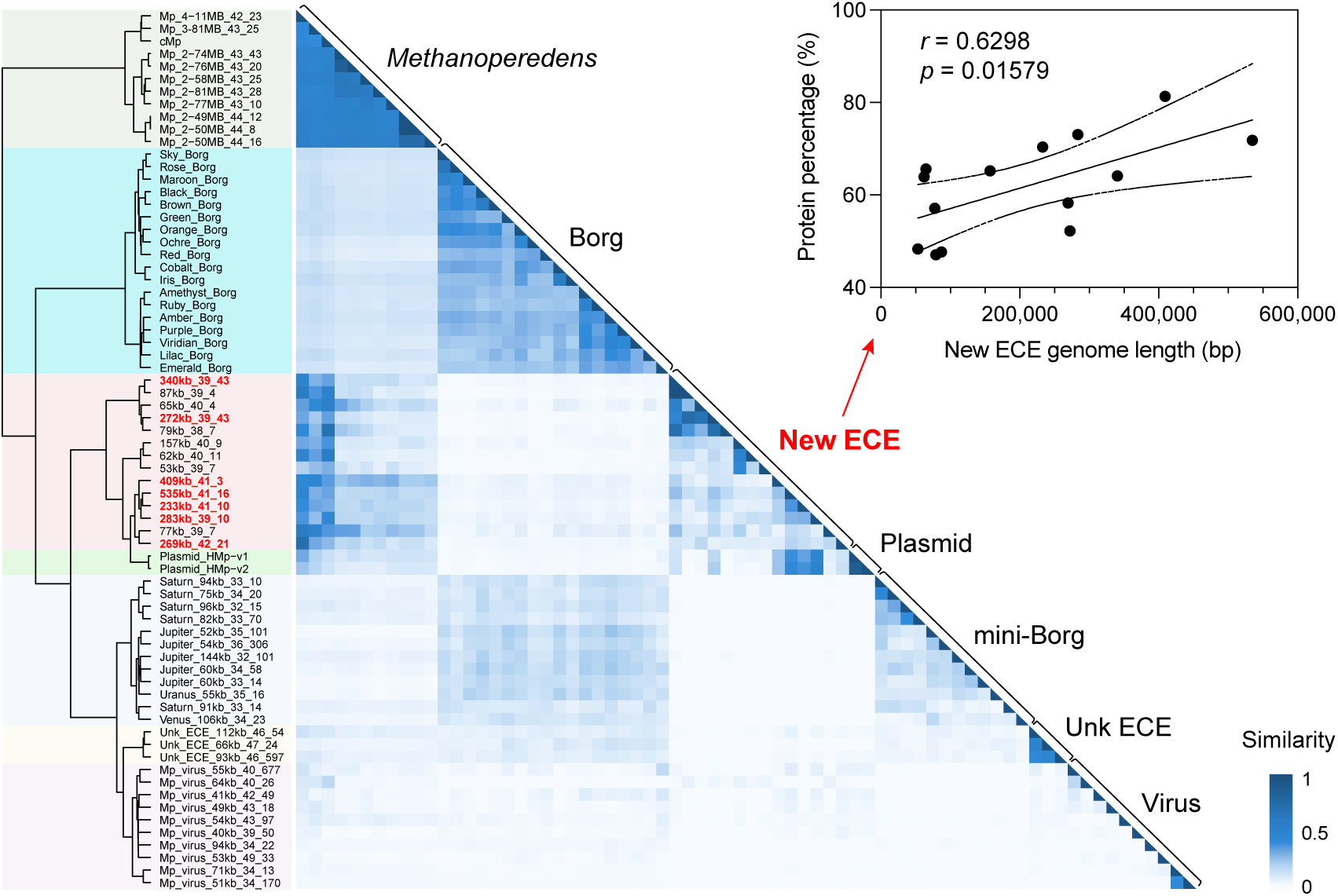
Comparison of new ECE genomes to *Methanoperedens* and the other associated ECEs. Genomes are clustered based on pairwise proteome similarity scores. Linear regression was performed on the new ECE genome length and the percentage of proteins that best match those in *Methanoperedens*. Dashed lines indicate errors at 95% confidence of the best-fit line. The regression coefficient *r* and *p*-value were calculated by Pearson correlation and two-sided t-test.

Given that organisms likely methylate their DNA to enable distinction between “self” and “not self”, it is hypothesized that ECEs that associate with hosts share specific host DNA methylation patterns. Modified bases can be detected using the nucleotide incorporation kinetics that are recorded during PacBio sequencing. We therefore searched for DNA modification motifs using PacBio HiFi reads and found consistent modification patterns between certain ECEs and *Methanoperedens* genomes (Supplementary Fig. 4). The results support the ECE-host linkages inferred from the high sequence identity of transposons (∼2 kb). Based on the DNA modification patterns and sequence similarity, Methanoperedens_40-02_00 and very closely related strains apparently host five of the ECEs, two of which have > 200 kbp genomes (Supplementary Figs. 3 and 4). Three of the ECEs are likely hosted by strains of Methanoperedens_43-46_00, and a Mp_4-11MB_42_23 strain is the likely host for one jumbo ECE. We calculated abundances of each ECE and the *Methanoperedens* inferred to be hosts and found ECE : host ratios of 1.44 ± 0.24 (Supplementary Table 3). Abundance ratios were constant in cases where the ECEs and inferred hosts co-occurred in two or more samples (Supplementary Fig. 5 and Supplementary Table 3).

We then compared the overall proteome similarity between the newly discovered ECEs, their host *Methanoperedens* archaea, and all previously described ECEs that associate with the same host (e.g., linear Borgs and mini-Borgs, circular viruses and plasmids). The proteomes of the new ECEs are distinct and clade together, with most similarity to the *Methanoperedens* proteomes (Fig. 1). Consistent with this, the ECE sequences have comparable GC contents to those of the *Methanoperedens* genomes (38.2 – 42.0% versus 40.6 – 44.1%) (Supplementary Tables 1 and 2). They do not encode any of the viral structural proteins (e.g., capsid) or plasmid hallmark genes (e.g., relaxase), according to annotations based on sequences and structures (Supplementary Table 4).

### New jumbo ECEs encode chromosome-type replication machinery

DNA replication origin recognition genes (Orc1) were identified in all jumbo ECEs (including the integrated one) and in some of the smaller ECEs (Supplementary Fig. 4). Most of these Orc1 proteins are phylogenetically close to those of *Methanoperedens*, but three affiliate closely with those of other archaea (e.g., Archaeoglobi) (Supplementary Fig. 4). Flanking the Orc1 proteins, all ECEs have AT-rich intergenic regions (e.g., 69.0 ± 6.3% of AT in the upstream 100 bp), and one (272kb_39_43) has short, direct tandem repeats, possibly iterons that may facilitate DNA unwinding and replication initiation.

Sequence- and structure-based annotations indicate that all of the new ECEs likely lack replicative helicases and thus must recruit host helicases during replication, as observed in other ECEs^30,31^. However, the jumbo ECEs encode numerous DNA repair-related helicases likely used to remove DNA lesions and maintain genome fidelity (Supplementary Table 4)^32^. Also encoded in most jumbo ECEs are archaeal, catalytic primase subunit (PriS, Supplementary Fig. 7), DNA polymerase B (PolB, Supplementary Fig. 8), and non-histone chromosomal protein (MC1, Supplementary Fig. 9). The ECE PriS proteins form two phylogenetic clades, both closely related to the *Methanoperedens* homologs. The proteins are structurally similar to the characterized PriS from the archaeon *Pyrococcus*, but lack a domain comprised of α-helices (Supplementary Fig. 7). This domain has an unknown function and is known to be highly variable in length and sequence among archaea^33^. The ECE PolB proteins are nested within the Pol B2 phylogenetic clade that comprises *Methanoperedens* and some Borg sequences (Supplementary Fig. 8). The ECE MC1 proteins, likely involved in genome organization, place phylogenetically with *Methanoperedens* sequences and have predicted structures that align well with the characterized MC1 protein from *Methanosarcina* (PDB: 2NBJ) (Supplementary Fig. 9). Overall, the new jumbo ECEs appear to encode *Methanoperedens*-like, chromosome-type replication machinery.

### Jumbo ECEs may expand *Methanoperedens* metabolic capacity

Jumbo ECEs encode genes that are absent in the host. One example is a protein with a ThiF family domain that occurs in, for instance, thiamine biosynthesis protein ThiF, ubiquitin activator Uba, and molybdopterin-synthase adenylyltransferase MoeB, all of which are structurally and functionally similar to each other^34^. The ThiF family proteins encoded by the jumbo ECEs are distantly related to the single-copy homologs in *Methanoperedens* (Fig. 2a) and have conserved residues at the active site (Fig. 2b). Interestingly, ThiS family proteins that would be activated by ThiF-like enzymes are present in two or three copies in each *Methanoperedens* genome (Fig. 2c). We predicted the structure of a heterodimeric complex comprising the ECE ThiF-like protein and the *Methanoperedens* ThiS-like protein and obtained confident interfaces between these protein chains (AlphaFold3 ipTM = 0.80, pTM = 0.86; Fig. 2d). Particularly, the C-terminal Gly-Gly tail of the ThiS-like protein extends into the pocket of the partner ThiF-like protein for activation and adenylation^35^. As *Methanoperedens* harbor fewer ThiF family proteins than ThiS counterparts, we infer that the ECE proteins most structurally similar to ThiF may complement the host protein pool and interact with the extra *Methanoperedens* ThiS-like proteins (Fig. 2).

**Figure 2.**
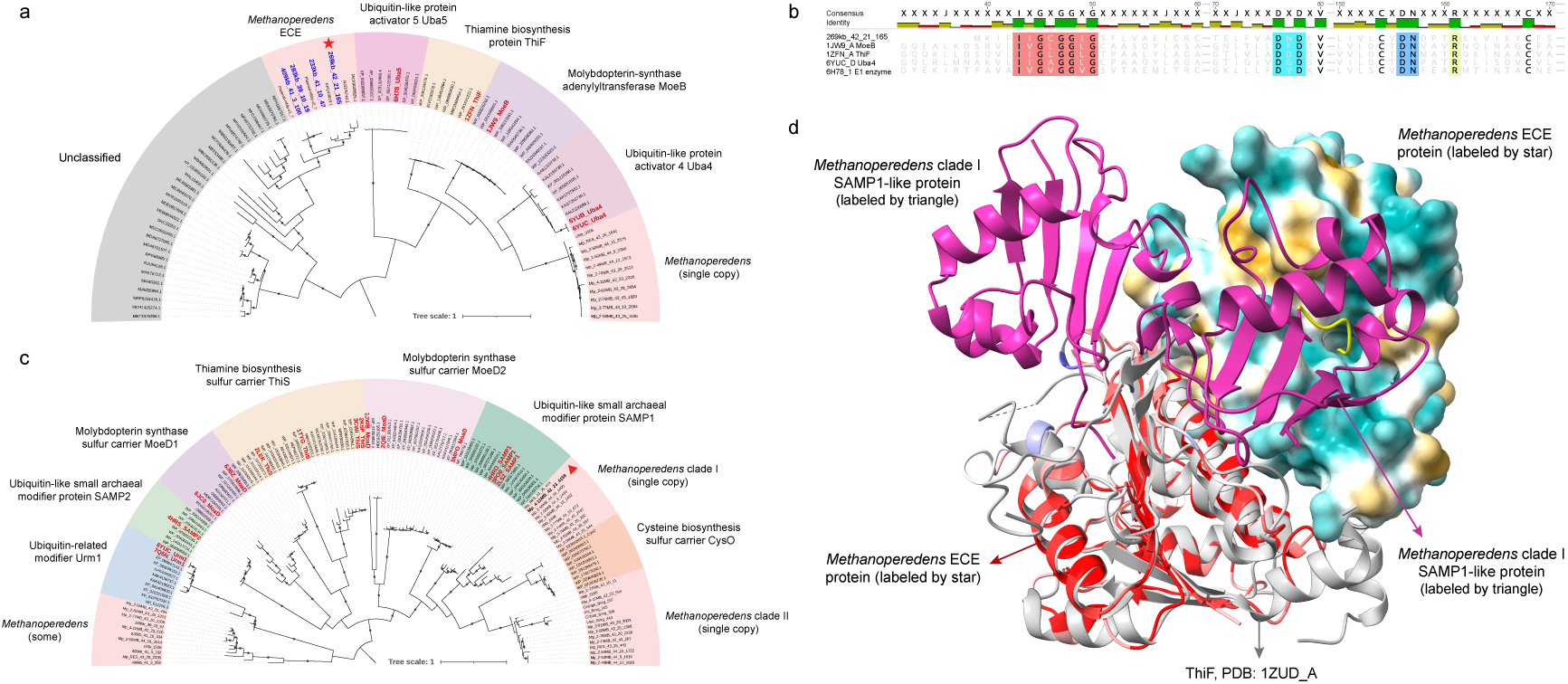
Sequence and structural analyses of ThiF and ThiS family proteins in *Methanoperedens* and jumbo ECEs. **(a)** Phylogeny of proteins containing the ThiF family domain. Sequences in bold blue are from jumbo ECEs, and those in bold red have experimentally resolved structures. Dots on branches indicate the same topology observed in > 80% of 1000 re-samplings. **(b)** Alignment of representative proteins in the ThiF family. The red box includes highly conserved residues (in black), which indicate a glycine-rich, nucleotide-binding motif proposed to facilitate ATP entry into the active site. The cyan box shows ribose-binding sites. The blue box is responsible for the proper positioning of Asp130, which may be involved in Mg^2+^ ligation. The yellow box is predicted to support the C-terminal Gly-Gly dipeptide of partner proteins (e.g., MoeD). **(c)** Phylogeny of proteins containing the ThiS family domain. Sequences in bold red have experimentally resolved structures. Dots on branches indicate the same topology observed in > 80% of 1000 re-samplings. **(d)** Heterodimer comprising the *Methanoperedens* SAMP1-like protein (labeled by triangle) and the jumbo ECE protein (labeled by star) folded by AlphaFold3. The resulting ipTM and pTM values are 0.80 and 0.86. One heteromonomer is colored based on the predicted local distance difference test (pLDDT) score. ThiF (PDB: 1ZUD_A; gray) is the best structural match of the ECE protein, and thus, they are superimposed together. C-terminal Ala-Val-Ser-Gly-Gly peptides of the *Methanoperedens* SAMP1-like protein are colored in yellow. One copy of the ECE protein is displayed in the surface mode and colored by lipophilicity using the default ChimeraX coloring scheme.

Jumbo ECEs encode genes that could be involved in anaerobic methane oxidation. A formylmethanofuran dehydrogenase subunit B is encoded in the 272kb_39_43 and is closely related to the protein in *Methanoperedens* (Supplementary Fig. 10a). In methanotrophic archaea, formylmethanofuran dehydrogenase catalyzes the conversion of formylmethanofuran to CO_2_, a final step in anaerobic oxidation of methane^36,37^. This protein complex has several different versions in *Methanoperedens*, all of which contain subunit B (Supplementary Fig. 10b). The ECE protein is likely a part of the formylmethanofuran dehydrogenase subunit DBAC, according to the sequence phylogeny (Supplementary Fig. 10a), and is structurally similar to the homolog from methanogenic *Methanothermobacter* (PDB: 5T5I) (Supplementary Fig. 8c). Also encoded in one jumbo ECE (409kb_41_3) are tetrahydromethanopterin S-methyltransferase (Mtr) subunits A, D, and E, parts of the eight-subunit complex that converts methyl-coenzyme M (CH_3_-S-CoM) to methyl-tetrahydromethanopterin (CH_3_-H_4_MPT)^38^, and F_420_H_2_:NADP^+^ oxidoreductases (FNO) that catalyze the interconversion of F_420_ and F_420_H_2_^39^, a key cofactor for anaerobic methane oxidation (Supplementary Table 4). These proteins all share high similarity with the homologs in *Methanoperedens*.

Jumbo ECEs encode genes related to nitrogen compound transformations. A copper-containing nitrite reductase that is never found in the host *Methanoperedens* is encoded by 233kb_41_10 and 283kb_39_10 (Supplementary Fig. 11). These ECE proteins are predicted to form a homotrimer that aligns well with the crystal structure of nitrite reductase from the anammox bacterium *Candidatus* Jettenia caeni (PDB: 5ZL1). Three jumbo ECEs encode Mo-nitrogenases falling into Group IV-A and Group VII^40^, both of which groups are also present in a complete *Methanoperedens* genome (Supplementary Fig. 12a). The nitrogenase gene operons from ECEs include at least NifHDK and are highly similar to operons in *Methanoperedens* (Supplementary Fig. 12b). The NifD and NifK proteins are predicted to form a heterodimeric complex, as expected based on the literature^41^ (Supplementary Fig. 12c-d). Many ECEs also encode molybdate/tungstate and other types of ABC transporters that transport metals across membranes for nitrogenase and other metalloenzymes (Supplementary Table 4).

One jumbo ECE (409kb_41_3) encodes a molybdopterin oxidoreductase that belongs to the dimethylsulfoxide reductase (DMSO) family^42^. Sequence phylogeny suggests that the DMSO family protein is a putative dissimilatory sulfur reductase subunit A (SreA) closely related to those identified in *Methanoperedens* in a prior study^43^ (Supplementary Fig. 13a). Downstream of the ECE *sreA* are genes encoding [4Fe-4S]-containing SreB and integral transmembrane SreC for electron transfer (Supplementary Fig. 13b). The ECE SreABC proteins are predicted to form a membrane-bound heterodimer towards the (pseudo-)periplasmic space, consistent with the structural configurations of many DMSO family proteins, e.g., the polysulfide reductase complex^44^ (Supplementary Fig. 13c). Upstream of the ECE *sreA* are genes highly similar to *arrR*, *arrS*, and *arrX* (Supplementary Fig. 13b), which are known to regulate the expression of the Arr complex, another member within the DMSO family^45^.

Jumbo ECEs encode genes predicted to be involved in other forms of energy conservation. One ECE 409kb_41_3 carries part of the F_420_H_2_ dehydrogenase (Fpo complex), a homolog of respiratory NADH-ubiquinone oxidoreductase (Nuo complex), which oxidizes F_420_H_2_ coupled to the translocation of protons^43^. The ECE transmembrane proteins FpoL, M, and N have predicted structures that align well with biochemically characterized structures of bacterial NuoLMN and are highly similar to the homologs from the host *Methanoperedens* in sequence (Supplementary Fig. 14). We therefore cofolded the ECE FpoLMN with the other components of the *Methanoperedens* Fpo complex and observed confident protein-protein interactions (Supplementary Fig. 14), suggesting that the jumbo ECE may augment host energy generation. Also encoded in ECE genomes are NDP-forming acetyl-CoA synthetase and pyruvate kinase that generate ATP along with the production of acetate and pyruvate, respectively (Supplementary Table 4). Another enzyme able to produce ATP is anaerobic carbon monoxide dehydrogenase (CODH), which may interconvert CO and CO_2_^46,47^. This enzyme is encoded in two jumbo ECEs (340kb_39_43 and 283kb_39_10, Supplementary Fig. 15a). The ECE proteins (blue) cluster phylogenetically apart from CO_2_-fixing CODH (in red) and CODH variants (in yellow) but have six conserved residues required to coordinate a Ni-Fe-S cluster (Supplementary Fig. 15b)^48^. They also show very similar structural conformation to an experimentally resolved CODH involved in CO oxidation (Supplementary Fig. 15c), and thus, we speculate that these ECE proteins likely perform irreversible CO oxidation. In addition, multiheme *c*-type cytochromes (MHCs) that enable electron transfer for energy conservation are prevalent in the ECEs, with up to 18 heme-binding motifs (Supplementary Table 4). Three jumbo ECEs encode the complex cytochrome *c* maturation (Ccm) system I, which can load hemes into apocytochromes for MHC maturation (Supplementary Fig. 16)^49^.

Some ECE genes form operons that catalyze independent, sequential reactions. An intriguing example is the ∼6 kbp genomic region identified in 272kb_39_43 that encodes two partial tRNA ligases, an archease that may enhance the function of tRNA ligase, and two transposases (Supplementary Fig. 17a). One transposase interrupts the tRNA ligase gene, and the other apparently perturbs the archaease gene, resulting in a disordered C-terminus (relative to *Pyrococcus*, PDB: 4N2P). A highly similar region occurs in the *Methanoperedens* genome 4-11MB_42_23, but without transposases (Supplementary Fig. 17b). We speculate that the ECE acquired the archease-RNA ligase operon from the host *Methanoperedens*, but it was subsequently inactivated by transposon insertion.

Another operon related to tRNA processing is the *tusBDCA*, which is part of the tRNA thiolation system (Numata et al., 2006), found in three jumbo ECEs, and a *Methanoperedens* genome proximal to a tRNA (Supplementary Fig. 18). The ECE TusD subunit has a conserved cysteine at the active site for sulfur transfer and is confidently predicted to form a heterohexameric complex with TusB and TusC (ipTM 0.93 and pTM 0.92), which are encoded adjacent to TusD (Supplementary Fig. 18). Also identified is a four-gene operon in 340kb_39_43 that encodes histidine ammonia-lyase, formiminoglutamase, urocanase, and imidazolonepropionase, all of which also occur in *Methanoperedens* and together catalyze a multi-step conversion from histidine to glutamate (Supplementary Fig. 19).

### Jumbo ECEs may extend *Methanoperedens* defense capability

ECEs encode genes for defense against invasive elements. A hybrid CRISPR-Cas system comprising Type I and Type III-A components is identified in 269kb_42_21 (Fig. 3a). None of the Cas proteins in the ECE have homologs in any of the four CRISPR-Cas systems of the organism predicted to be the host based on transposon sequence similarity (Mp_4-11MB_42_23, Supplementary Table 4). However, Cas6 in the ECE locus has homologs in the putative host genome, but not within the CRISPR-Cas regions (Fig. 3b).

**Figure 3.**
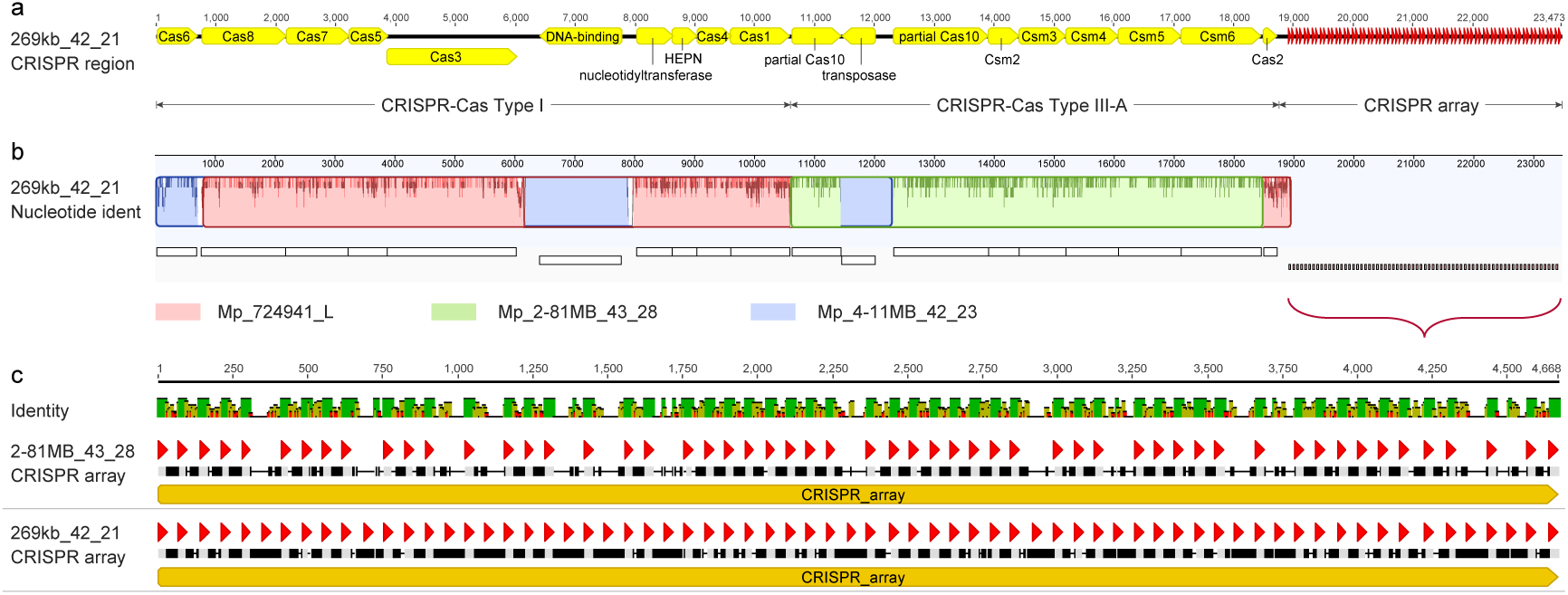
A hybrid CRISPR-Cas system in the jumbo ECE acquired from multiple *Methanoperedens* species. **(a)** Genomic context of the hybrid CRISPR-Cas system in the jumbo ECE 269kb_42_21. Yellow arrows indicate predicted genes, and red ones indicate interspaced 30-bp repeats. **(b)** Mauve alignment of the hybrid CRISPR-Cas system. Colored chunks along the ECE genome show the best blastn matches from different *Methanoperedens* genomes and corresponding nucleotide similarities. **(c)** MAFFT alignment of CRISPR arrays in the *Methanoperedens* 2-81MB_43_28 and the jumbo ECE 269kb_42_21. Gray vertical bars indicate identical sequences, and black bars indicate different ones. Red arrows demonstrate the 30-bp repeats (SNP ≤ 1) in the 2-81MB_43_28 (n = 55) and the 269kb_42_21 (n = 69).

A region highly similar to the Type I part of the Type I/Type III-A system of ECE 269kb_42_21 occurs in *Methanoperedens* 724941_L, and a region similar to the Type III-A portion occurs in *Methanoperedens* 2-81MB_43_28 (Fig. 3b). These findings suggest a chimeric origin of the ECE CRISPR-Cas locus, and perhaps imply that 269kb_42_21 was resident in hosts of both types at some point. Given the similar Type III-A region and identical CRISPR repeats in the ECE and *Methanoperedens* 2-81MB_43_28, we infer that *Methanoperedens* 2-81MB_43_28 is the likely host for the 269kb_42_21. Notably, the ECE shares no CRISPR spacers with the possible host (Fig. 3c), so it may augment the host defense capacity.

Other genes implicated in host defense (and possibly ECE-ECE competition) include partial Type III-A CRISPR-Cas (Csm) systems that include only Csm3, Csm4, and short CRISPR arrays (Supplementary Fig. 20). One such region is encoded in 340kb_39_43 and two in 79kb_38_7. In addition, there are non-specific targeting, Type I restriction-modification systems comprised of HsdR, HsdM, HsdS, and YhcG, identified in 340kb_39_43 and 269kb_42_21, with very similar regions in *Methanoperedens* (Supplementary Fig. 21).

ECEs also encode genes implicated in extracellular competition. Interestingly, the 269kb_42_21 genome encodes a system structurally similar to an extracellular contractile injection system (eCIS) that may deliver effector proteins such as toxins into target cells or the surrounding environment (Supplementary Fig. 22)^50^. Similar genomic regions are also found in *Methanoperedens*. A protein structurally comparable to Cagβ-like protein from *Helicobacter pylori* (PDB: 8DOL) and potentially part of a Type IV system capable of secreting toxins is encoded in 272kb_39_43 but not in *Methanoperedens* (Supplementary Fig. 23)^51^. The putative ECE Cagβ-like protein is confidently predicted to form a homohexamer (iPTM 0.73 and pTM 0.78) with an ∼3.5 nm central channel and a membrane anchor (Supplementary Fig. 23C and D). Sequence-based phylogenetic analysis suggests a bacterial origin of this protein. The ECE genomes also encode diverse toxin-antitoxin (TA) systems (Supplementary Fig. 24). In most cases, more toxins were identified than antitoxins, and the expected, paired antitoxins of some TA systems were not found.

### Expression of genes encoded in jumbo ECEs

We obtained metatranscriptomic data from a wetland sediment column that contained two jumbo ECEs and determined that they were actively transcribing metabolic genes. Highly expressed proteins in 283kb_39_10 included the cobalt transporter complex CbiMOQ, which mediates the uptake of cobalt essential for Mtr function, nitrite reductase (NirK), carbon monoxide dehydrogenase (CODH), and the nitrogenase components NifH and NifK (Fig. 4a; Supplementary Table 5). Also expressed are replication-related proteins, such as Orc1 and PriS.

**Figure 4.**
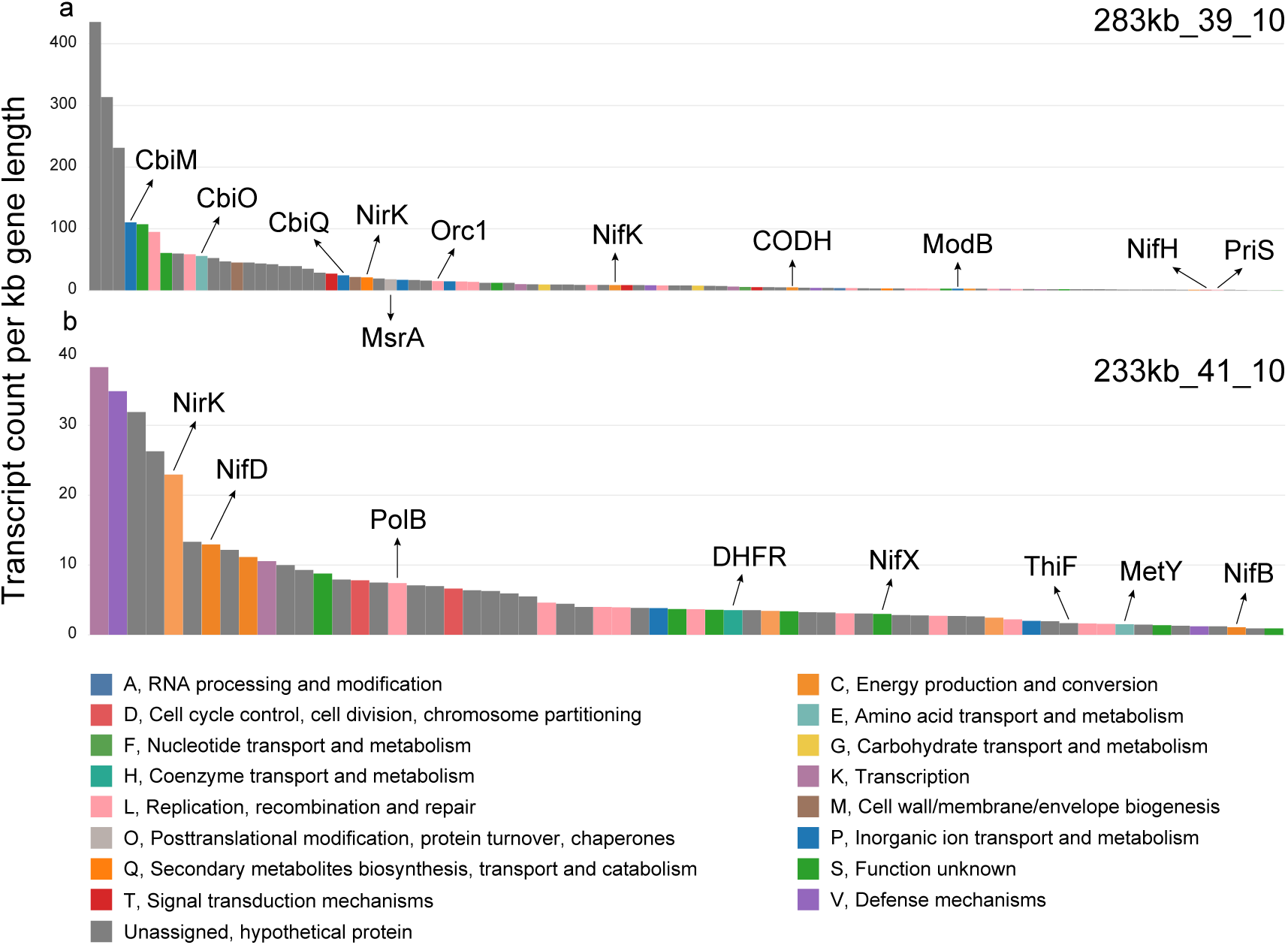
Gene expression profile of jumbo ECEs. *In situ* activity of 283kb_39_10 **(a)** and 233kb_41_10 **(b)** in a wetland sediment column. Genes are shown as bars in the x-axis and colored based on COG category. The y-axis indicates the expression of each gene in the count of transcript read pairs divided by the gene length in kilobases. Protein abbreviations: CbiM, cobalt transporter transmembrane protein; CbiO, cobalt transporter ATP-binding protein; CbiQ, cobalt transporter associated protein; NirK, copper-containing nitrite reductase; MsrA, methionine sulfoxide reductase; Orc1, DNA replication origin recognition protein; NifK, Mo-nitrogenase beta chain; CODH, anaerobic carbon monoxide dehydrogenase; ModB, molybdate/tungstate transporter, permease protein; NifH, nitrogenase iron protein; PriS, catalytic primase subunit; NifD, Mo-nitrogenase alpha chain; PolB, DNA polymerase B; DHFR, dihydrofolate reductase; NifX, FeMo cofactor biosynthesis protein; ThiF, thiamin biosynthesis protein; MetY, O-acetyl-L-homoserine aminocarboxypropyltransferase; NifB, FeMo cofactor biosynthesis protein.

Gene expression by 233kb_41_10 genes was one order of magnitude lower than for 283kb_39_10 (Fig. 4). However, nitrite reductase, nitrogenase NifD and its accessory proteins NifBX, and thiamin biosynthesis protein ThiF were relatively highly expressed (Fig. 4b; Supplementary Table 5). Also active were DNA polymerase B and dihydrofolate reductase, both of which contribute to the ECE genome replication.

### Genome expansion of jumbo ECEs

Given the large number of ECE genes that are highly similar to those in *Methanoperedens*, we speculated that jumbo ECEs underwent genome expansion by acquiring DNA material from hosts. Thus, we compared the nucleotide sequences of the *Methanoperedens* and ECE genomes. As jumbo ECEs encode core metabolic genes, it is difficult to confidently distinguish ECE from *Methanoperedens* genome fragments. To overcome this, we reconstructed 10 new, circular, complete *Methanoperedens* genomes from the sites where ECEs were discovered. These, and a previously reported complete *Methanoperedens* genome from the wetland site^52^, enabled accurate ECE and *Methanoperedens* sequence comparisons. The jumbo ECEs share many DNA blocks (> 500 bp) of high similarity (> 80% nt identity) with each other. Genomic regions in larger ECEs (e.g., 535kb_41_16 and 409kb_41_3) without related sequences in smaller ECEs always have highly similar counterparts in *Methanoperedens*, suggestive of many DNA acquisition events from host *Methanoperedens*. Notably, the jumbo ECEs also share many DNA blocks with a putative plasmid (HMp-v2, ∼155 kbp) previously described from a *Methanoperedens* bioreactor enrichment^27^ (Fig. 5). Unlike the jumbo ECEs, the HMp-v2 plasmid contains few nucleotide regions that are only related to those in *Methanoperedens* (i.e., there are related sequences in other ECEs; Fig. 5).

**Figure 5.**
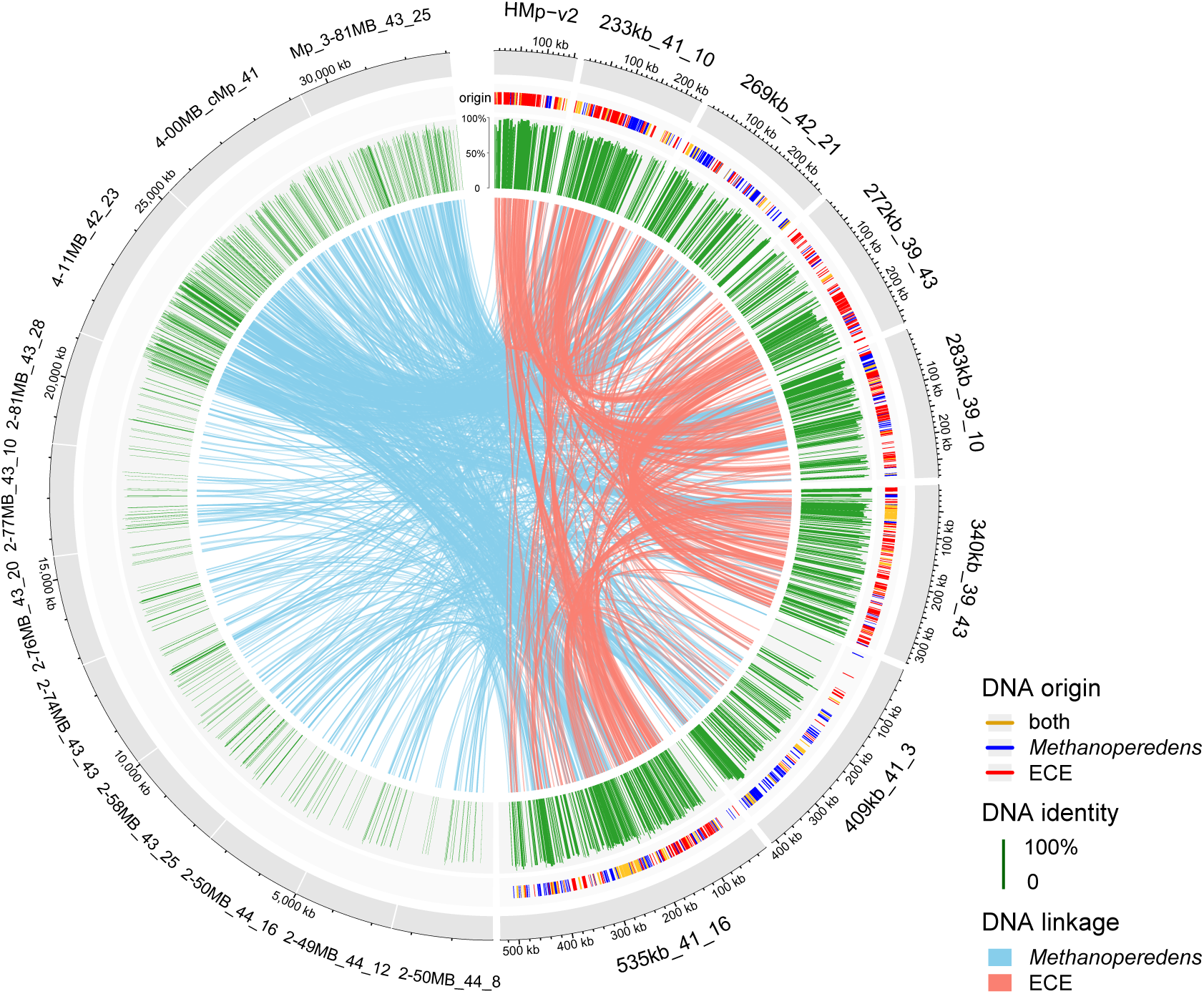
Expansion of ECE genomes. 11 circular *Methanoperedens* genomes are placed in the left half of the outer ring; seven jumbo ECEs discovered here and one previously reported plasmid are on the right. Note that the scales of *Methanoperedens* and ECE genomes are different, for visualization purposes. The second outer ring indicates that the ECE regions are shared with *Methanoperedens* only (blue), or with ECEs only (red), or with both (gold). The length of green bars in the third ring shows the DNA identity of shared genomic regions. Inner chords link shared blocks across different genomes. Only blocks that have lengths > 500 bp and pairwise identities > 80% are depicted and connected.

## Discussion

Of all genetic elements, archaeal ECEs with large genomes are the least well characterized. Here, we report seven novel, jumbo ECEs for which we reconstructed complete genomes of up to 535 kbp in length (and an additional seven complete ECEs < 200 kbp). Based on protein sequence similarity, we infer that these ECEs all associate with anaerobic methane-oxidizing *Methanoperedens* archaea. Notably, one 409 kbp element is integrated in a *Methanoperedens* genome, clearly establishing the host relationship. This integrated ECE represents the largest integrative element in the Archaea domain so far.

Unlike viruses and plasmids, the new ECEs encode *Methanoperedens*-like DNA processing genes that enable them to replicate in the same manner as the host. This distinguishes the ECEs from chromids, which utilize plasmid replication and partitioning systems^53^. Given the lack of certain proteins required for replication, they must recruit host machinery and thus remain under the host cell cycle control^54,55^. This may explain the relatively stable genome copy ratios under different conditions (1.44 ± 0.24) (Supplementary Fig. 3).

The jumbo ECEs provide genes not encoded in host genomes. These include defense systems such as the hybrid Type I/Type III-A CRISPR-Cas systems, likely derived from different *Methanoperedens* species (Fig. 3). Together with other CRISPR-Cas and restriction-modification systems (Figs. S18 and S19), the ECE genes may reinforce protection of the host against invading viruses, plasmids, and other elements (Fig. 6). They also encode unpaired, orphan toxins (Supplementary Fig. 24). Toxins and antitoxins in plasmids work interactively to ensure plasmid survival and transmission by killing daughter cells that lose these elements^56,57^. Given that ECEs carry eCIS and Type IV secretory systems, we postulate that they may secrete and/or inject these toxins into surrounding organisms, thus reducing extracellular competition^50^ (Fig. 6).

**Figure 6.**
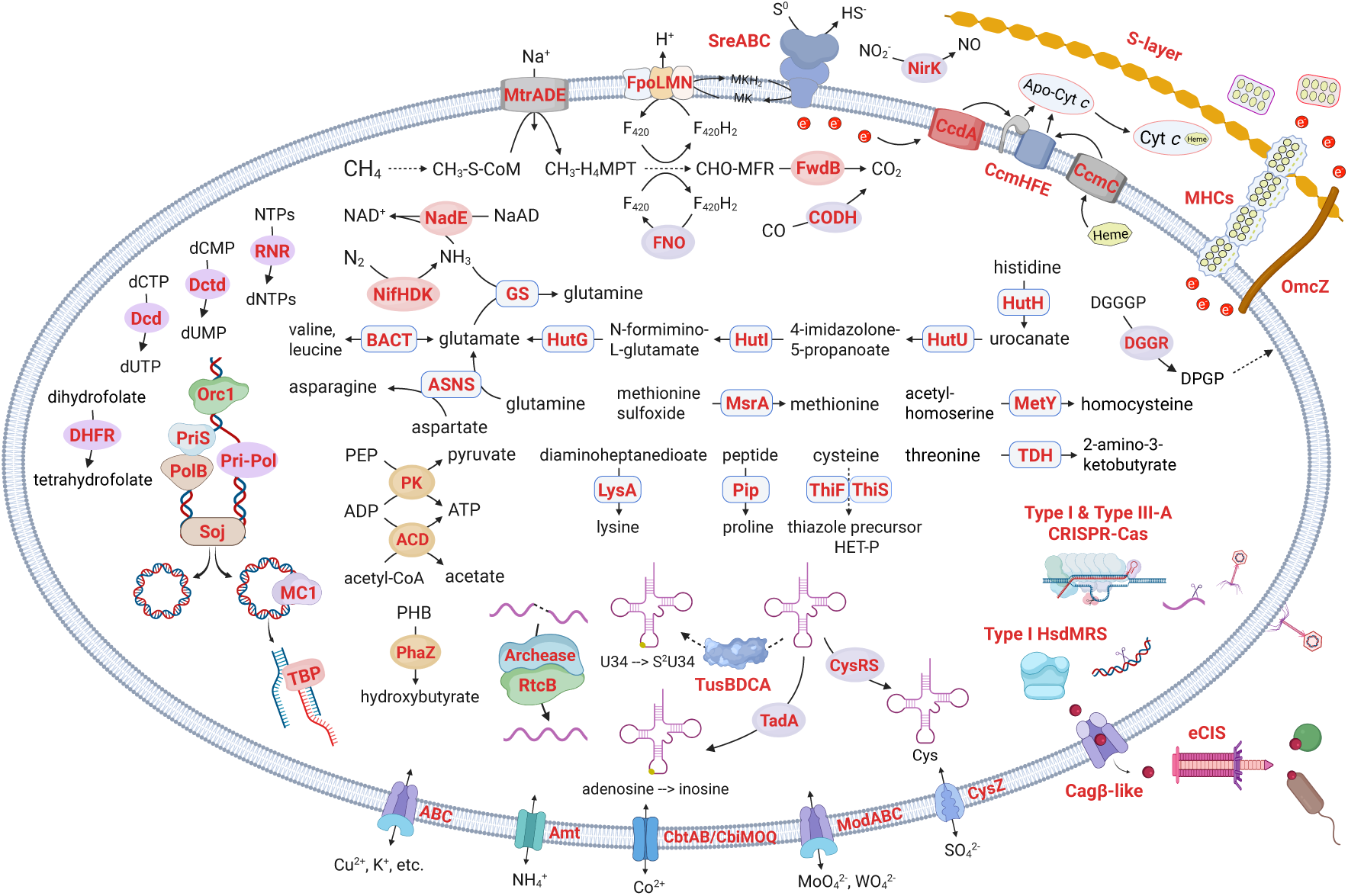
Function of jumbo ECEs in *Methanoperedens* cells. Proteins encoded in jumbo ECEs are highlighted in red and bold. Mtr, tetrahydromethanopterin S-methyltransferase; Fpo, F_420_H_2_ dehydrogenase; Sre, sulfur reductase; NirK, copper-containing nitrite reductase; CcdA, thiol:disulfide interchange protein; Ccm, cytochrome maturation system; MHCs, multiheme cytochromes; OmcZ, nanowire-form multiheme cytochrome; Fwd, formylmethanofuran dehydrogenase; CODH, anaerobic carbon monoxide dehydrogenase; FNO, F_420_H_2_:NADP^+^ oxidoreductase; NadE, NAD synthetase; Nif, Mo-nitrogenase; GS, glutamine synthetase; RNR, anaerobic ribonucleotide reductase; Dctd, dCMP deaminase; Dcd, dCTP deaminase; DHFR, dihydrofolate reductase; Orc1, DNA replication origin recognition protein; PriS, catalytic primase subunit; PolB, DNA polymerase B; Pri-Pol, bifunctional DNA primase/polymerase; Soj, chromosome partitioning ATPase; MC1, archaeal chromosomal protein; TBP, TATA binding protein; BACT, branched-chain amino acid aminotransferase; HutG, N-formylglutamate deformylase; HutI, imidazolonepropionase; HutU, urocanate hydratase; HutH, histidine ammonia-lyase; ASNS, asparagine synthetase; MsrA, methionine sulfoxide reductase; MetY, O-acetyl-L-homoserine aminocarboxypropyltransferase; TDH, threonine dehydrogenase; DGGR, digeranylgeranylglycerophospholipid reductase; PK, pyruvate kinase; ACD, NDP-forming acetyl-CoA synthetase; PhaZ, PHB depolymerase; LysA, diaminopimelate decarboxylase; Pip, proline iminopeptidase; Thi, thiamin biosynthesis protein; RtcB, RNA ligase; Tus, tRNA-specific sulfur transferase; TadA, tRNA-specific adenosine deaminase; CysRS, cysteinyl-tRNA synthetase; Hsd, restriction-modification system; Amt, ammonium transporter; Cbt/Cbi, cobalt transporter; Mod, molybdate/tungstate transporter; CysZ, sulfate permease protein; Cagβ, Cagβ-like secretory protein; eCIS, extracellular contractile injection system. Detailed gene annotation is present in Supplementary Table 3.

Notably, the jumbo ECEs encode dissimilatory sulfur reductase (not encoded in the *Methanoperedens* genomes reported here), nitrite reductase (expressed, and absent in *Methanoperedens*), and multiheme cytochromes. Multiheme cytochromes are also encoded by *Methanoperedens*, but the jumbo ECE sequence variants likely differ in their redox potentials (Dolla et al., 1994), thus could expand the host’s ability to pass electrons to extracellular mineral electron acceptors. The ECE-specific sulfur and nitrite reductase and multiheme cytochromes may increase the host’s physiological plasticity by linking either CH_4_ or CO oxidation to multiple electron sinks. CO oxidation by *Methanoperedens* is indicated by expression of ECE-encoded anaerobic carbon monoxide dehydrogenase (Fig. 4) (also encoded by the host). Two recent studies report that *Methanoperedens* (formerly ANME-2d) and other ANME-2 lineages can oxidize CO when CH_4_ is absent^58,59^.

Most functional genes in ECEs are also present in hosts with high similarities. (Fig. 6). Conspicuous is tetrahydromethanopterin S-methyltransferase, an enzyme known to function only in methane metabolism, catalyzing the second step of anaerobic methane oxidation. Also involved in this pathway are ECE-associated formylmethanofuran dehydrogenase and F_420_H_2_:NADP^+^oxidoreductase, which together enable ECEs to augment the host methane-oxidizing capacity. Other examples of genes present in ECEs and hosts include genes for nitrogen fixation, tRNA modification, and amino acid metabolism (Fig. 6).

An intriguing question is why ECEs carry sequence variants of genes also present in their host organism’s genomes. One explanation is that ECEs genes increase gene copy number, thus augmenting specific metabolic activities. Alternatively, the gene variants may expand metabolic capacity due to different substrate specificities and/or reaction rates. Some variants may be resistant to targeted disruption caused by stressors such as phage defense systems or antibiotics^60^, thus increasing the fitness of hosts in the environment.

The large inventory of metabolic genes is reminiscent of Borgs, the recently identified ECEs of *Methanoperedens* with up to 1.1 Mbp linear genomes^24^. However, the jumbo ECEs described here have fundamentally different genome architecture – they are circular. They have higher GC contents of ∼40% than Borgs of ∼33% and more genes with high sequence similarity to those of *Methanoperedens* (∼62% versus ∼20%).

Multiple genome alignments clearly show genomic relatedness among the jumbo ECEs, the host *Methanoperedens*, and a previously reported plasmid (Fig. 5), raising the possibility of evolutionary relatedness. Perhaps jumbo ECEs evolved from fragments of host chromosomes^61^, as they contain numerous genomic blocks highly similar to those in *Methanoperedens* genomes (Fig. 5). Gene loss may explain the evolution to relatively small plasmids. Alternatively, small plasmids may have persistently incorporated DNA from hosts and evolve into jumbo ECEs, as genomic regions that are present in jumbo ECEs but not in plasmids always occur in *Methanoperedens* (Fig. 5). Given that jumbo ECEs have chromosome-type replication machinery and encode large inventories of metabolic genes, we conclude that they fall on a spectrum between minichromosomes and plasmids, revealing a continuum in the evolution of different ECE types associated with *Methanoperedens* archaea.

## Methods

### DNA extraction, sequencing, and processing

Soil samples were collected at depths ranging from 80 to 110 cm below the surface at a wetland site in Lake County, California, USA. One sample of 20 - 30 cm deep was collected at a rice field in Butte County, California, USA. Approximately 5.0 g of soil per sample was used for DNA extraction with the Qiagen DNeasy PowerMax Soil Kit. Extracted DNA was sequenced using the PacBio HiFi long-read technology and the Illumina short-read platform.

Low-quality raw reads were trimmed and/or removed using BBDuk (https://jgi.doe.gov/data-and-tools/software-tools/bbtools/). Qualified long reads were assembled into scaffolds using hifiasm-meta (v0.13-r308)^62^ and metaMDBG (v0.3)^63^. Short reads were assembled using metaSPAdes (v4.2.0)^64^. Scaffolds longer than 1 kbp were binned using a combination of CONCOCT (v1.1.0)^65^, MaxBin2 (v2.2.7)^66^, MetaBAT2 (v2.15)^67^, and VAMB (v3.0.2)^68^. Generated bin sets were dereplicated and optimized using DAS Tool (v1.1.2)^69^.

### Identification and curation of archaeal ECE genomes

Genes in assembled scaffolds were predicted by Prodigal (v2.6.3) using the DNA code 11^70^. Taxonomy was assigned to each protein using “mmseqs easy-taxonomy” (version: 6f45232ac8daca14e354ae320a4359056ec524c2)^71^ based on the best hit in the GTDB v226^72^. Putative “self-circular” genomes produced by PacBio HiFi assemblies were classified using GTDB-Tk (v2.3.0) against the prokaryotic reference database (v226)^73^. Genomes confidently assigned to taxonomy were considered microbial. We initially focused on the unclassified, putatively circular genomes that are > 200 kbp in length, lack rRNAs and ribosome components, and have proteins with best matches to those of Archaea, as typical of archaeal jumbo ECEs.

PacBio HiFi reads were mapped to candidate ECE genomes using Minimap2 (v2.28-r1209) in the “map-hifi” mode with default parameters^74^. Illumina reads were mapped using BBMap (https://github.com/BioInfoTools/BBMap). The generated SAM files were converted to BAM formats by SAMtools (v1.17)^75^ and visualized in Geneious Prime (2025.0.3). For “self-circular”, jumbo ECE genomes from PacBio HiFi assemblies, long reads mapping was used to verify circularization, followed by curation using short reads mapping to ensure base call support at every position and remove assembly errors. The validated ECE genomes were then used to identify related genome fragments, including those < 200 kbp sequences. Reads were mapped to recruited scaffolds using the same method as described above. Those placed at scaffold ends and the unplaced paired Illumina reads were used to extend the sequences. In some cases, the extended sequences can join with other scaffolds. This process was repeated until the further extension in one end of the genome came back to the other end, indicative of circularization. Details of curation can be found in our previous paper^76^.

### Curation of *Methanoperedens* genomes

Using the same method above, PacBio long reads and Illumina short reads were mapped to putative “self-circular” genomes that were classified as “f Methanoperedenaceae” in the GTDB taxonomy. We used read mapping to remove assembly errors and ensure base call consistency at every position in Geneious Prime (2025.0.3).

### Gene annotation based on sequences and structures

Proteins were compared with references in the KEGG, UniRef, and UniProt databases by USEARCH (v10)^77^, and in the NCBI nr database by BLASTp (v2.16.0+)^78^. Protein domains were predicted by HMMER3 (v3.3)^79^ against Pfam 37.0 using the pre-defined Noise Cutoff (NC) bit score thresholds^80^. COG category was assigned to proteins using eggNOG-mapper^81^. Toxins and antitoxins were identified and classified using TAfinder 2.0 on the TADB 3.0 web server^82^. To reduce false positives, we only retained hits that had either BLAST or HMMER e-values < 10^-5^. Subcellular localizations of proteins were predicted using PSORT (v3.0)^83^.

Proteins encoded by ECE genomes were structurally modeled using AlphaFold2 via LocalColabFold with default parameters^84,85^. Predicted structures labeled ‘relaxed_rank_001’ were compared with experimentally resolved structures in the PDB database using Foldseek (version 427df8a6b5d0ef78bee0f98cd3e6faaca18f172d)^86^. Protein functions were inferred from best-matching structural homologs meeting thresholds of *e*-values ≤ 0.001, bit scores ≥ 60, and query and target coverages ≥ 70%. Sequence- and structure-based annotations were integrated and cross-validated for each ECE protein.

### Genome comparison of *Methanoperedens* and associated ECEs

Pairwise similarity scores for complete *Methanoperedens* and associated ECE genomes were calculated based on global proteomes using DiGAlign^87^. Genomes were clustered and visualized using ‘ComplexHeatmap’ with Euclidean distance and Ward’s minimum variance method in R^88^.

### Genome linkage of individual ECEs to specific *Methanoperedens*

Linkages between ECE genomes and specific *Methanoperedens* genomes were predicted, in part, based on shared identical or near-identical transposon sequences, indicative of recent transfer of DNA material and thus the physical association of organisms^89^. In addition, DNA modification was inferred from PacBio sequencing data by leveraging polymerase kinetics, specifically inter-pulse duration (IPD) measured across reads mapped to each genome. For each genomic position, we estimated the expected (baseline) IPD with a k-mer–specific model learned directly from the metagenomic data, enabling control-free normalization of observed IPD values. Positions with significantly elevated IPD ratios were classified as modified, and modified sites were subsequently used for de novo motif discovery with MotifMaker (SMRT Link v13.1.0.221970). Abundances of ECE and *Methanoperedens* genomes were calculated from all available short-read datasets derived from the same site using CoverM v0.6.1^90^. As these genomes are comprised of single scaffolds, we used the ‘contig’ mode with a minimum read identity of 97% and a minimum aligned percent of 80% and kept trimmed mean coverages as abundances only if covered bases accounted for ≥ 50% of the references.

### Phylogenetic and structural analyses of ECE genes

Genes of interest were blasted against the NCBI nr database to recruit homologs. Both query and reference sequences were aligned using MAFFT (v7.453)^91^, followed by alignment trimming with trimAl (v1.4.rev15)^92^. Gene phylogeny was inferred based on sequence alignments using IQ-TREE (v2.3.6) with automatically selected best-fit models^93^. Phylogenetic trees were further decorated on the iTOL webserver^94^.

Structures of monomeric proteins were predicted mostly using AlphaFold2. Protein multimers were structurally modeled on the AlphaFold3 web server^95^. Structural analyses, including reference comparisons, surface electrostatic potential calculations, and channel size measurements, were executed in ChimeraX (1.7.1)^96^.

### RNA extraction, sequencing, and processing

Samples were collected from a sediment column that was inoculated with wetland soils and incubated in the lab. RNA was extracted using Qiagen RNeasy PowerSoil Total RNA Kit. Ribosomal RNA (rRNA) was depleted using RiboCop rRNA Depletion Kit (Lexogen) and QIAseq FastSelect – 5S/16S/23S Kit (Qiagen). The remaining RNA was used for library prep and sequenced on the Illumina NextSeq 2000 PE150 platform in QB3 Genomics, UC Berkeley.

Raw 150-bp paired reads were trimmed and/or removed using BBDuk. Quality-controlled reads were mapped to jumbo ECE genomes using BBMap with a minimum identity of 97%. Generated bam files were used to calculate mapped transcript reads for each gene using featureCounts (v2.0.6)^97^. Counts of read pairs were divided by gene length in kilobases for normalization.

### Multiple genome alignment of ECEs and *Methanoperedens*

Pairwise genome alignment among seven jumbo ECEs, 11 *Methanoperedens* genomes, and one previously reported plasmid was performed using MUMmer (v4.0.0rc1) with default parameters^98^. Blocks with length > 500 bp and nucleotide identity > 80% were retained and plotted using shinyCircos-V2.0 with modifications and decorations^99^. The *Methanoperedens* genome 3-81MB_43_25 was generated by removing the integrated jumbo ECE 409kb_41_3.

## Data availability

Metagenomic and metatranscriptomic sequencing reads of *Methanoperedens* and associated ECEs have been deposited in NCBI Sequence Read Archive (SRA) with the BioProject accession number: PRJNA1404836. Prior to publication, genomes can be accessed at: https://ggkbase.berkeley.edu/complete_Mp_HMp-plasmid-like/organisms.

## Acknowledgments

The authors would like to thank Tomas Hessler, Leylen Miloslavich, and Kaden DiMarco for their assistance in lab experiments. We thank LinXing Chen for the thoughtful comments on the manuscript. We thank Pacific Biosciences (PacBio) and the sequencing team in Menlo Park, CA, for providing HiFi sequencing for this project at no cost. We also thank Siyuan Zhang for technical support.

Funding for this research was provided by the Emerson Collective, the U.S. Department of Energy (DOE) Chemical Sciences, Geosciences, and Biosciences Division under Contract DE-AC02-05CH11231, and the Bill and Melinda Gates 901 Foundation (grant number INV-037174 to J.F.B.). The findings and conclusions are those of the authors and do not necessarily reflect positions or policies of the Bill and Melinda Gates Foundation. L.E.V.A. is supported by a Human Frontier Science Program Long-Term Fellowship (LT0043/2025-L). R.S. was funded by Lyda Hill Philanthropies, Acton Family Giving, the Valhalla Foundation, Hastings/Quillin Fund, the CH Foundation, Laura and Gary Lauder and Family, the Sea Grape Foundation, the Emerson Collective, Mike Schroepfer and Erin Hoffman Family Fund, and the Anne Wojcicki Foundation.

## Author Contributions

The study was designed by L-D.S. and J.F.B. L-D.S., B.K., L.E.V.A., and J.F.B. collected the soil samples. L-D.S., B.K., and L.E.V.A. extracted the DNA and assembled metagenomes. L-D.S. and J.F.B. identified and curated ECE and *Methanoperedens* genomes. L-D.S. performed genome and proteome analyses. L-D.S. extracted RNA and analyzed the metatranscriptomic data. S.W. and R.S. developed and used the PacBio methylation analysis methods. S.L. assisted in bioinformatics. L-D.S. and J.F.B. wrote the manuscript with input from all the authors.

