## Supplementary Figure for "Jumbo circular extrachromosomal elements of methane-oxidizing archaea with variably extensive metabolic and defense gene repertoires"

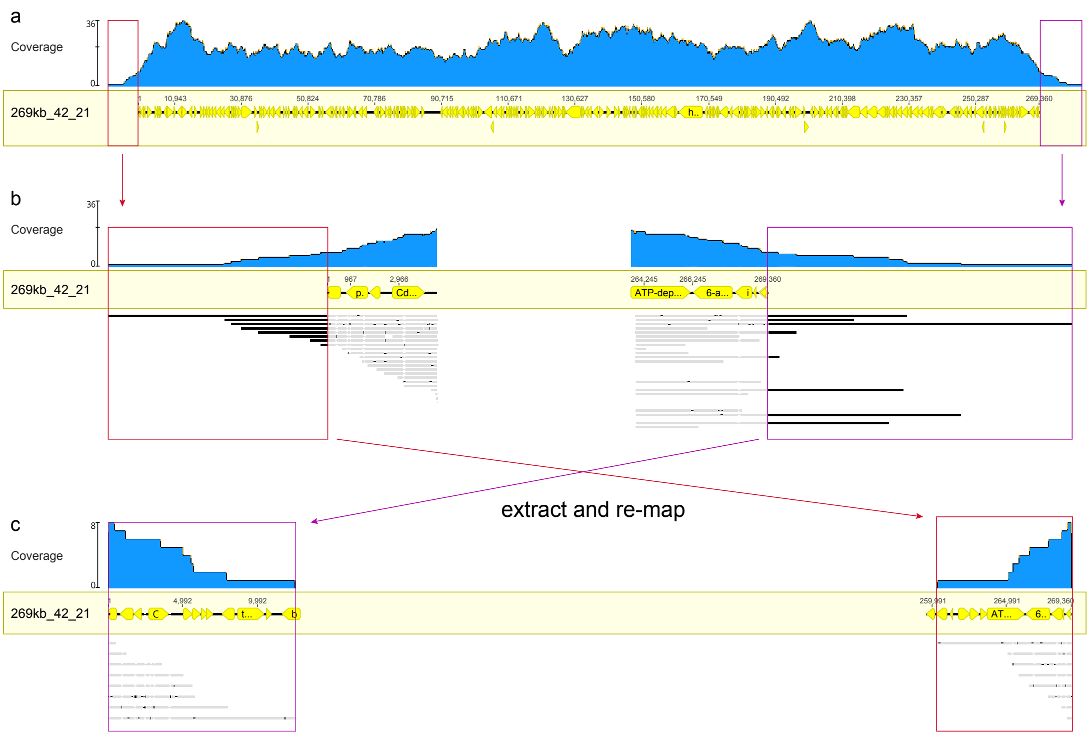


**Supplementary Figure 1. Validation of ECE genome circularization using PacBio long reads.** The genome 269kb_42_21 is used as an example showing the PacBio read mapping **(a)**. Extended read parts beyond the genome ends are highlighted and detailed in **(b)**. The extended parts were extracted and remapped to the genome to test the circularization in **(c)**. All the mapping was performed with mismatches per read ≤ 1%.

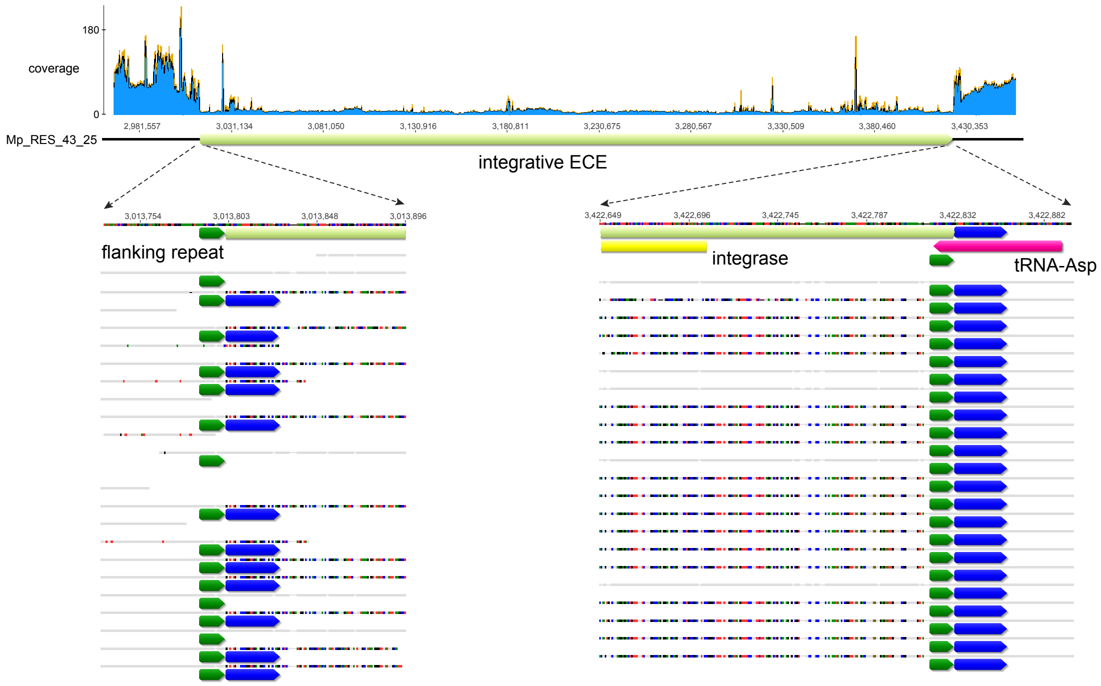


**Supplementary Figure 2. Integrative ECE in a *Methanoperedens* genome.** The mapping of PacBio and Illumina reads, allowing 2% mismatches per read, demonstrates a coverage dip in a genomic region. To capture partially mismatched reads, 30% of mismatches were allowed for a re-mapping, in which the ends of the integrative element are shown in detail. Gray bars below the genome are reads that match the reference. Colored parts inside reads are segments that do not match the reference. Arrows with the same colors indicate identical sequences. The partial, yellow arrow below the genome indicates the integrase. The pink arrow is the tRNA-Asp gene.

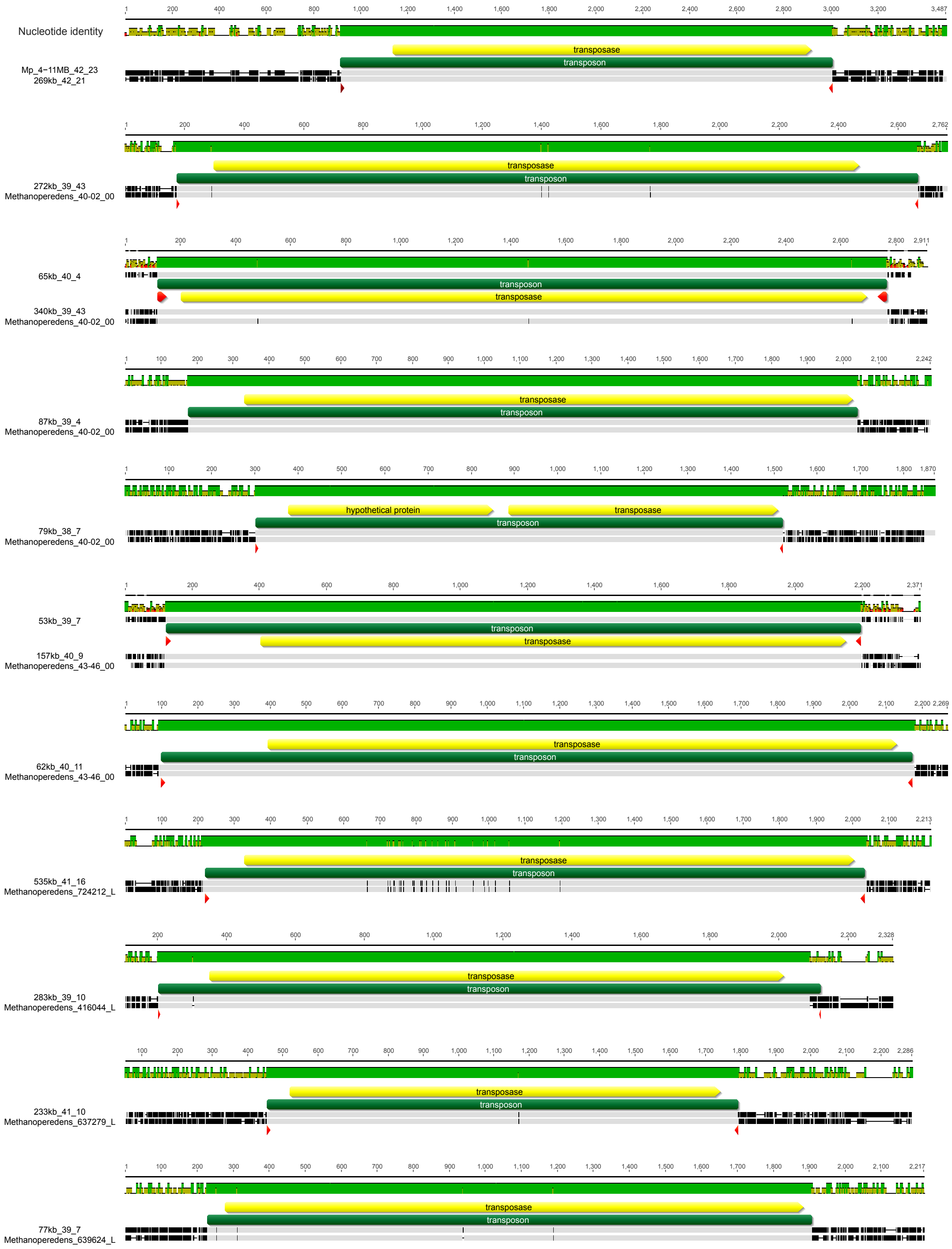


**Supplementary Figure 3. Transposon transfer between ECEs and *Methanoperedens*.** Genomic regions are aligned using Mafft. Gray bars indicate identical nucleotides and black bars indicate different ones. Green rectangles show transposons, in which yellow arrows indicate predicted transposases and red arrows indicate inverted repeats.

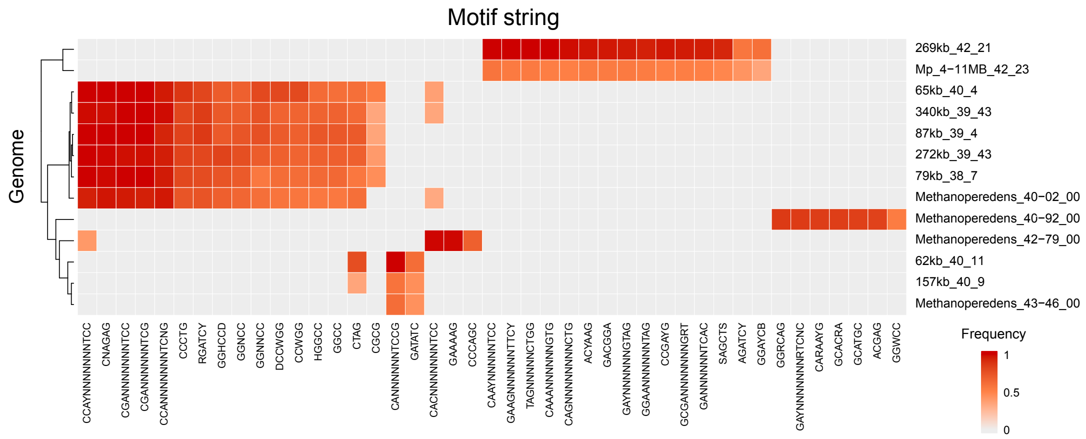


**Supplementary Figure 4. DNA methylation patterns of ECE and *Methanoperedens*.** Modified motifs were predicted for genomes that had sufficient abundances in a 100-cm deep sample using PacBio HiFi reads. Colors indicate the frequency of motifs that were methylated. Genomes are clustered based on Euclidean distances.

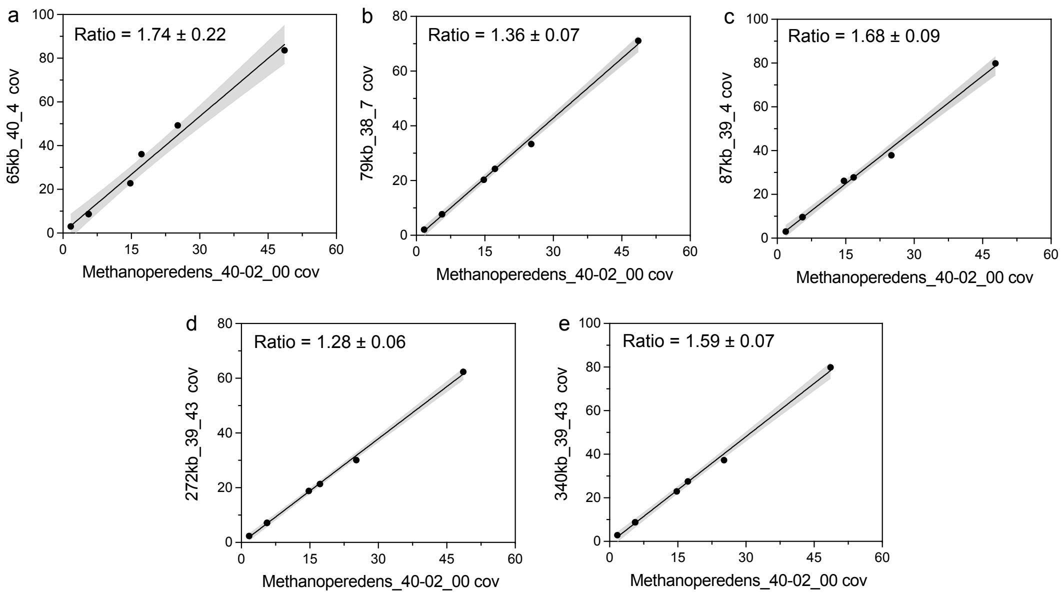


**Supplementary Figure 5. Co-occurrence of ECEs and predicted *Methanoperedens* species in multiple samples.** Scatters indicate pairwise genome abundances in the same sample and are fit by linear regression. Gray bands indicate errors at 95% confidence of the best-fit line. Average values and standard deviations were calculated for ECE-host ratios. ECEs are plotted only when they exist in more than three samples. Details and other ECE information are presented in Supplementary Table 3.

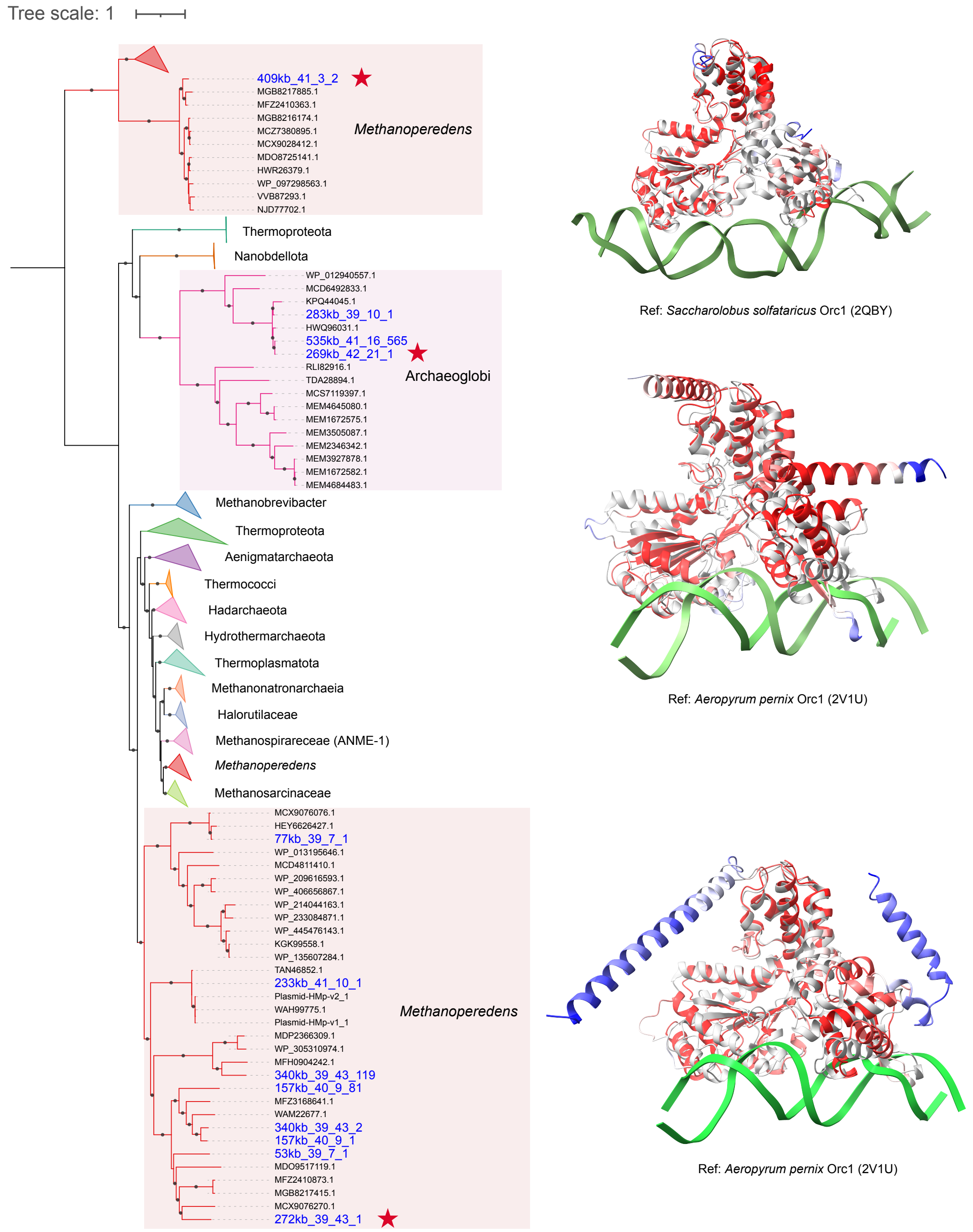


**Supplementary Figure 6. Phylogenetic and structural analyses of DNA replication origin recognition protein (Orc1).** The sequences labeled in blue are from the ECEs discovered in this study. Dots on branches indicate the same topology observed in > 80% of 1000 re-samplings. Structures of representative ECE sequences (labeled by stars) of each clade are predicted and superimposed with the best matching references that are experimentally resolved. The query structures are colored based on the pLDDT score. The reference protein structures are colored in gray and the DNA strands are in green.

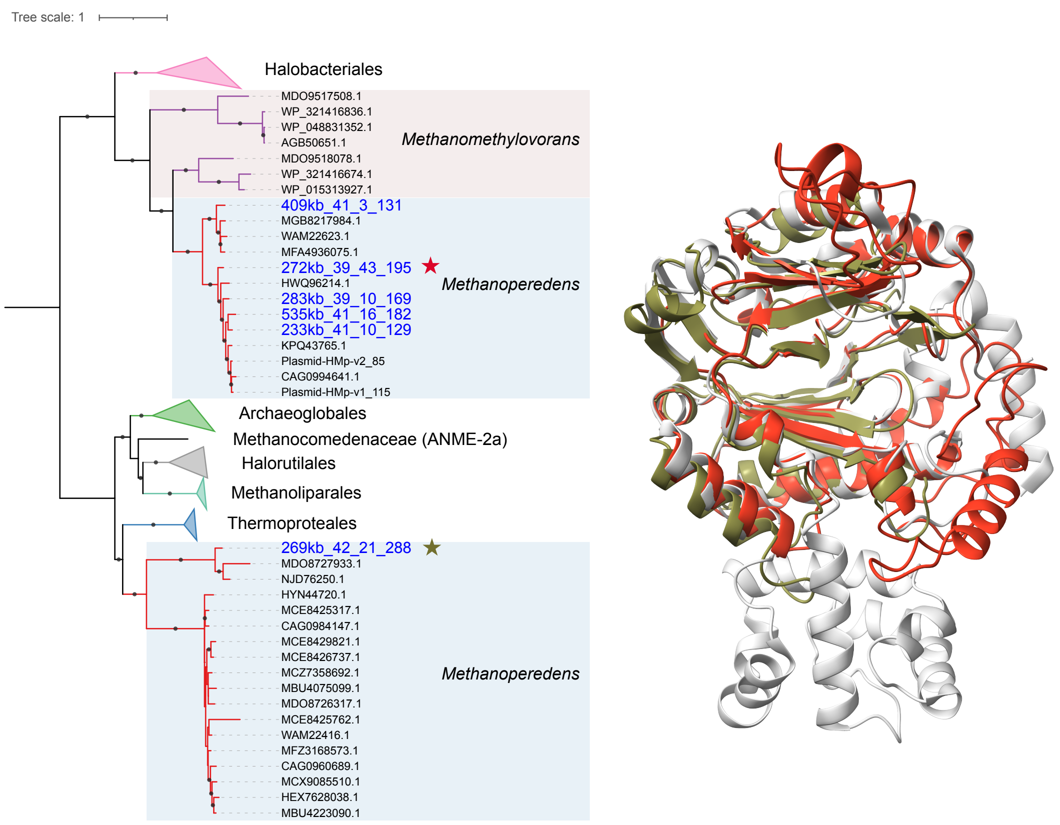


**Supplementary Figure 7. Phylogenetic and structural analyses of catalytic primase subunit (PriS).** Sequences in blue are from the ECEs discovered in this study. Dots on branches indicate the same topology observed in > 80% of 1000 re-samplings. Structures of the proteins labeled with stars are predicted, colored in red and green accordingly, and superimposed with the best matching reference in gray (PDB: 1G71). The extra domain in the reference structure but absent in queries has an unknown function and is highly variable in length and sequence within archaea and thus may be lost during evolution (https://www.nature.com/articles/nsb0101_57).

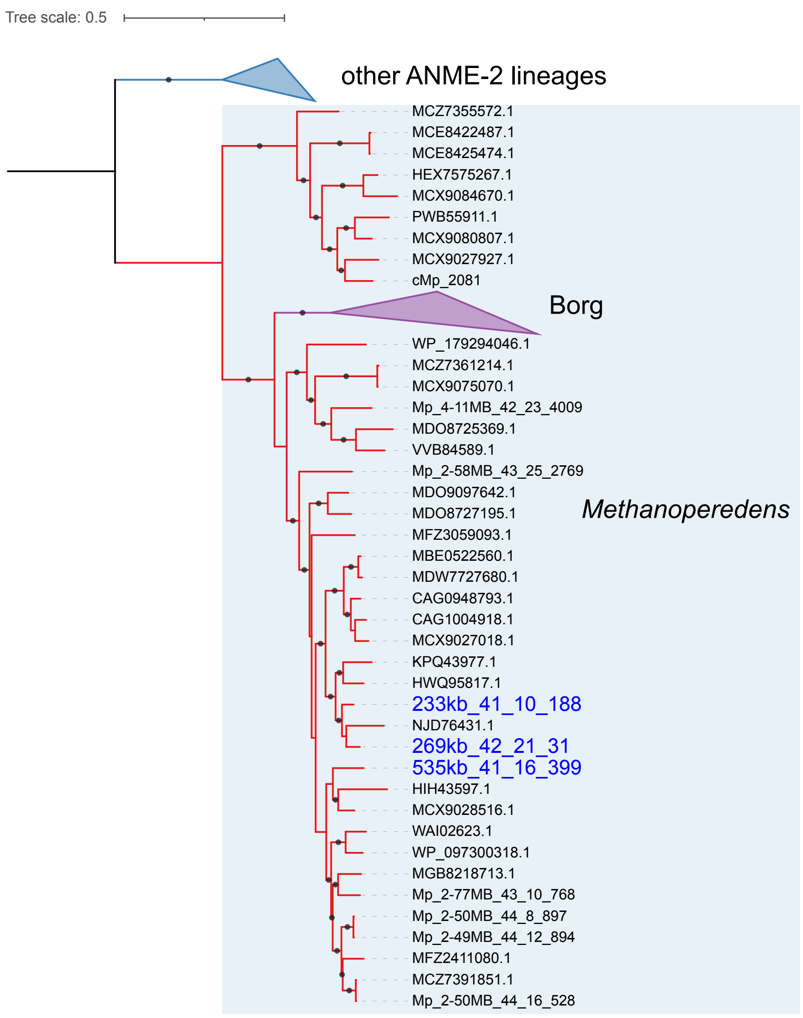


**Supplementary Figure 8. Phylogenetic analysis of DNA polymerase B encoded in jumbo ECEs.** The ECE sequences are labeled in blue. Dots on branches indicate the same topology observed in > 80% of 1000 re-samplings.

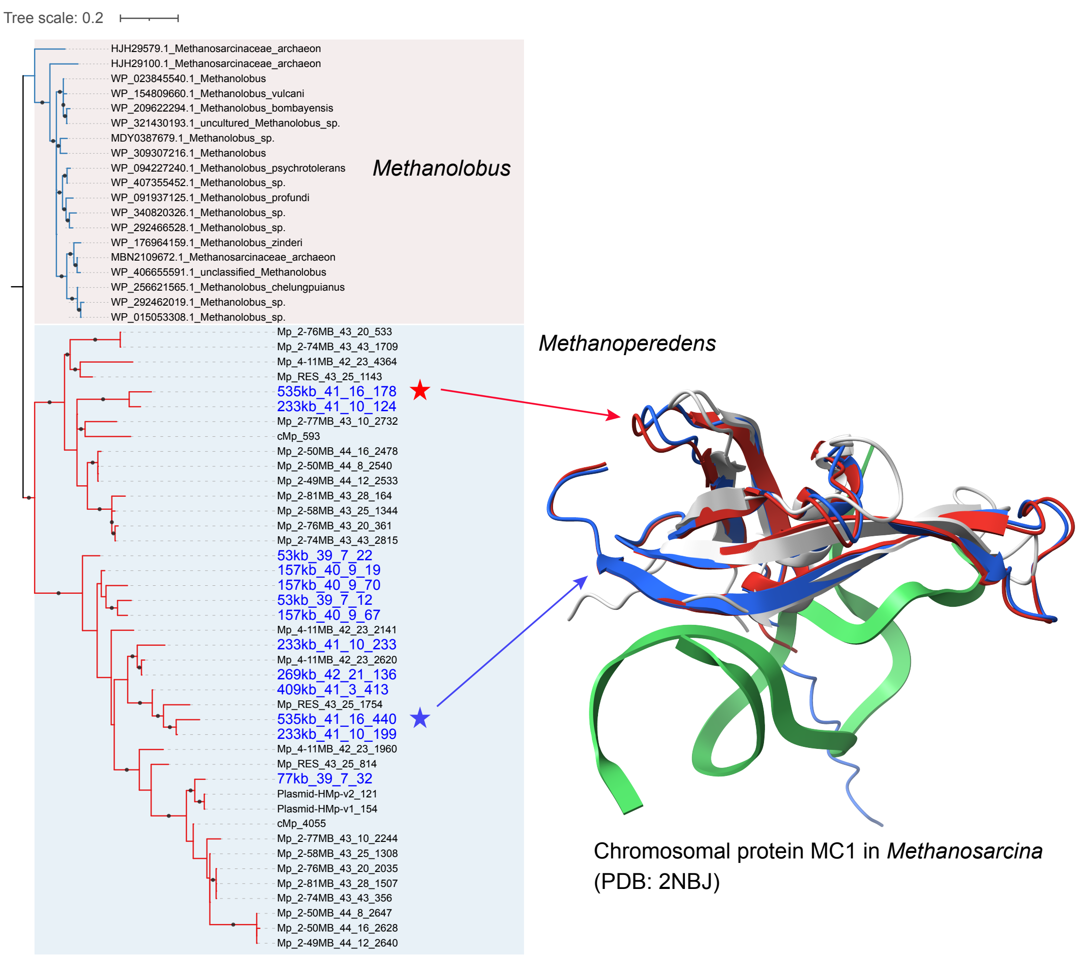


**Supplementary Figure 9. Phylogenetic and structural analyses of chromosomal protein MC1.** The proteins highlighted in blue are from the jumbo ECEs and smaller related ones. Dots on branches indicate the same topology observed in > 80% of 1000 re-samplings. Structures of the representative sequences labeled by stars are predicted and colored accordingly, and superimposed with the best matching reference in gray (PDB: 2NBJ).

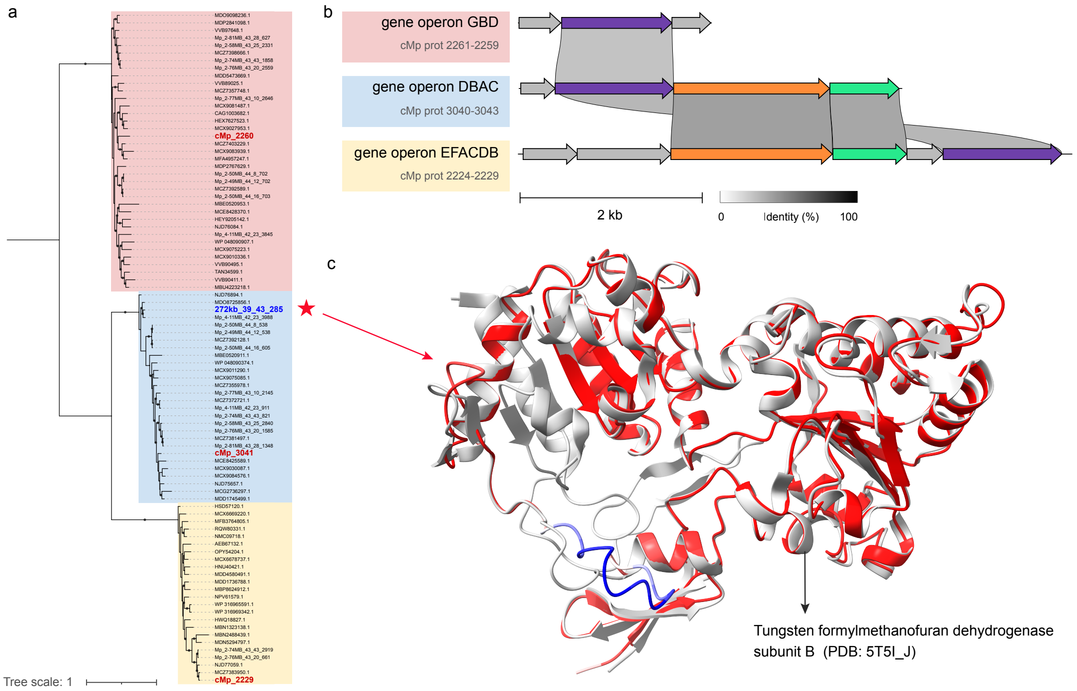


**Supplementary Figure 10. Sequence and structure analysis of formylmethanofuran dehydrogenase subunit B (FwdB). (a)** Phylogeny of formylmethanofuran dehydrogenase subunit B in *Methanoperedens* and the jumbo ECE 272kb_39_43. The ECE sequence is labeled a star. Dots on branches indicate the same topology observed in > 80% of 1000 re-samplings. **(b)** Three types of gene clusters containing formylmethanofuran dehydrogenase subunit B in the same, complete *Methanoperedens* genome (cMp). Colored shades of gene operon names correspond to sequence locations that are highlighted in red and bold in **(a)**. **(c)** Structure prediction and superimposition of formylmethanofuran dehydrogenase subunit B from 272kb_39_43. The query structure is colored based on the pLDDT score.

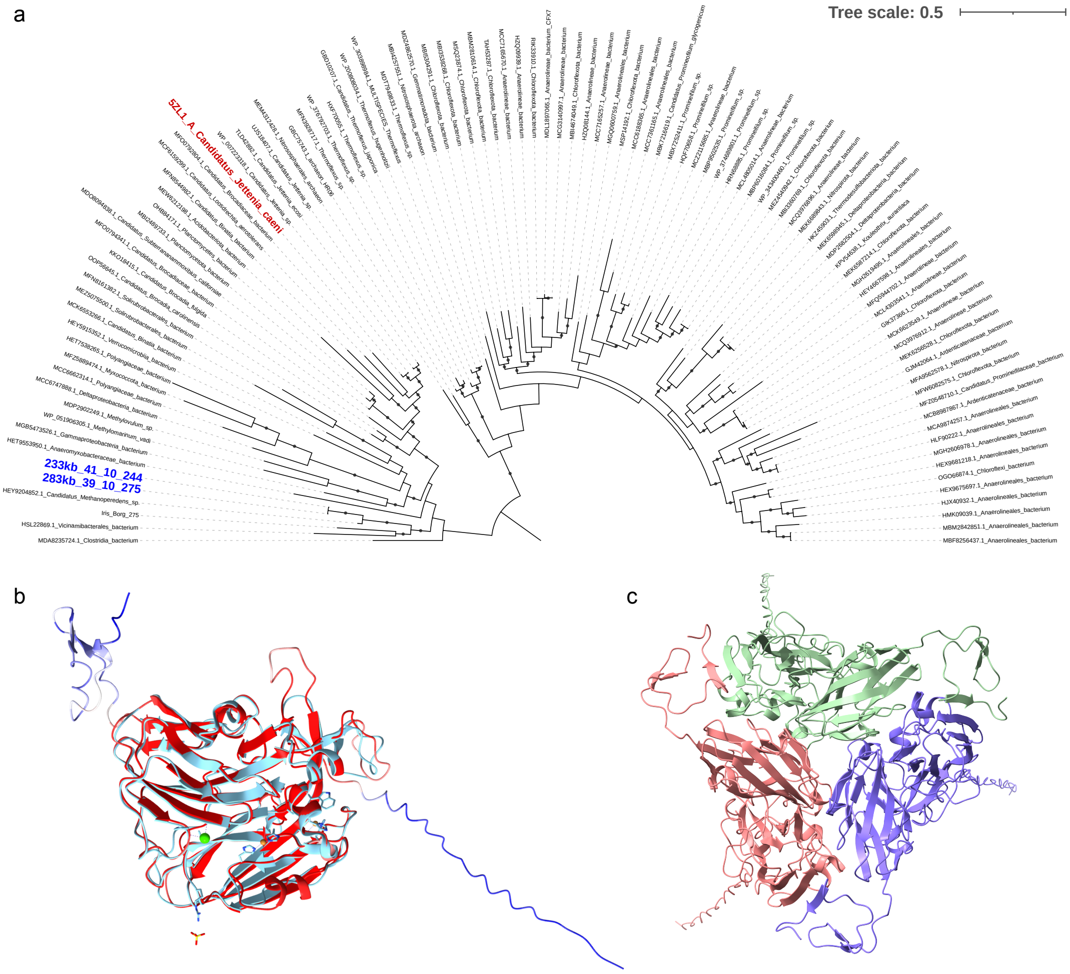


**Supplementary Figure 11. Copper-containing nitrite reductase encoded by jumbo ECEs. (a)** Phylogeny of copper-containing nitrite reductases. Proteins encoded by jumbo ECEs are highlighted in blue. The one in red has an experimentally resolved structure. Bootstrap values were calculated based on 1000 replicates and labeled as greater than 80%. **(b)** Structural superimposition of the ECE nitrite reductase (233kb_41_10_244) and the reference (5ZL1). The ECE protein structure is colored based on the pLDDT score. (**c**) Predicted homotrimer of the ECE nitrite reductase colored by chains. The ipTM and pTM values are 0.88 and 0.89.

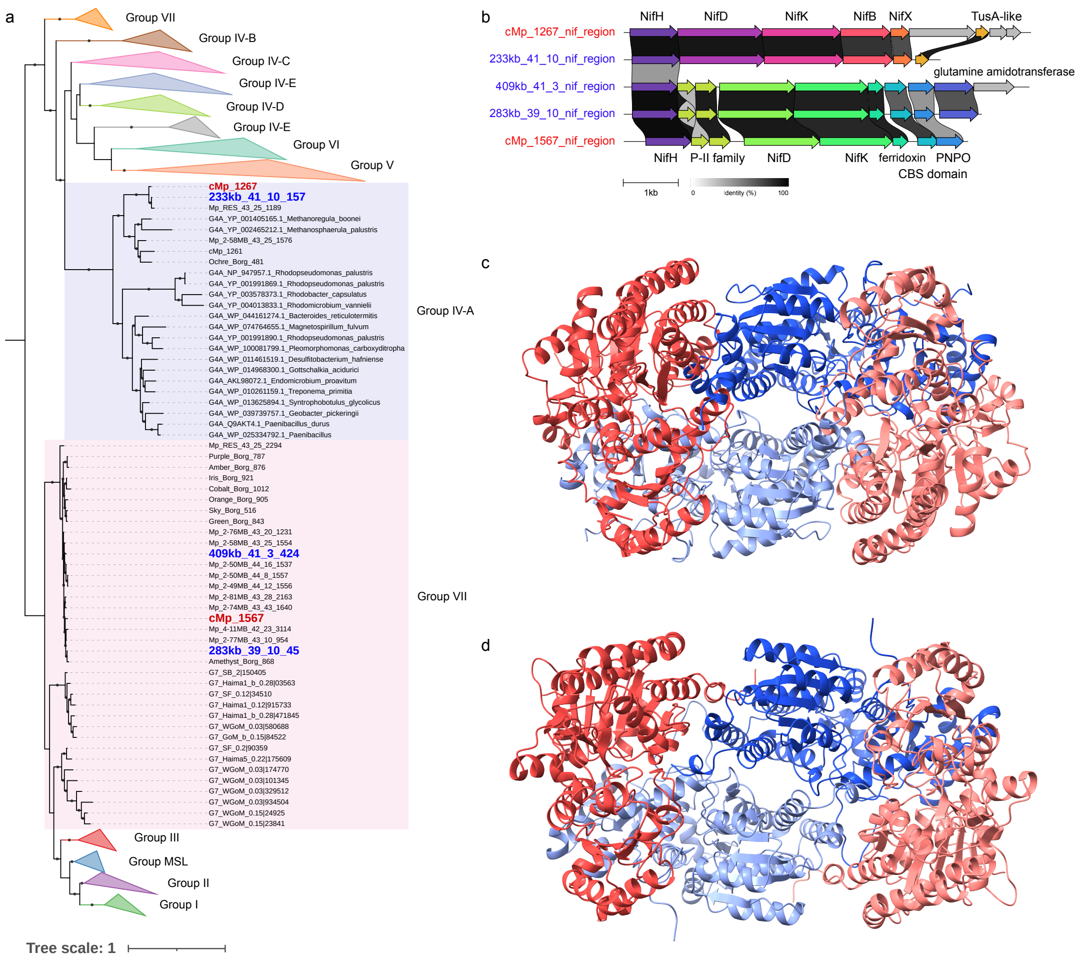


**Supplementary Figure 12. Nitrogenase encoded in jumbo ECEs. (a)** Phylogeny of NifH proteins. Sequences found in ECEs are highlighted in blue. Bootstrap values were calculated based on 1000 replicates and labeled as greater than 80%. **(b)** Comparison of gene clusters from ECEs and *Methanoperedens*. Homologous genes are connected based on pairwise identity > 30%. **(c-d)** Predicted heterodimeric structures of NifDK complexes encoded in jumbo ECEs. Alpha chains (NifD) are labeled in red-ish and beta chains (NifK) are in blue-ish. The ipTM and pTM values of the 233kb_41_10 complex are 0.92 and 0.93 **(c)**, and of the 409kb_41_3 complex are 0.94 and 0.95 **(d)**.

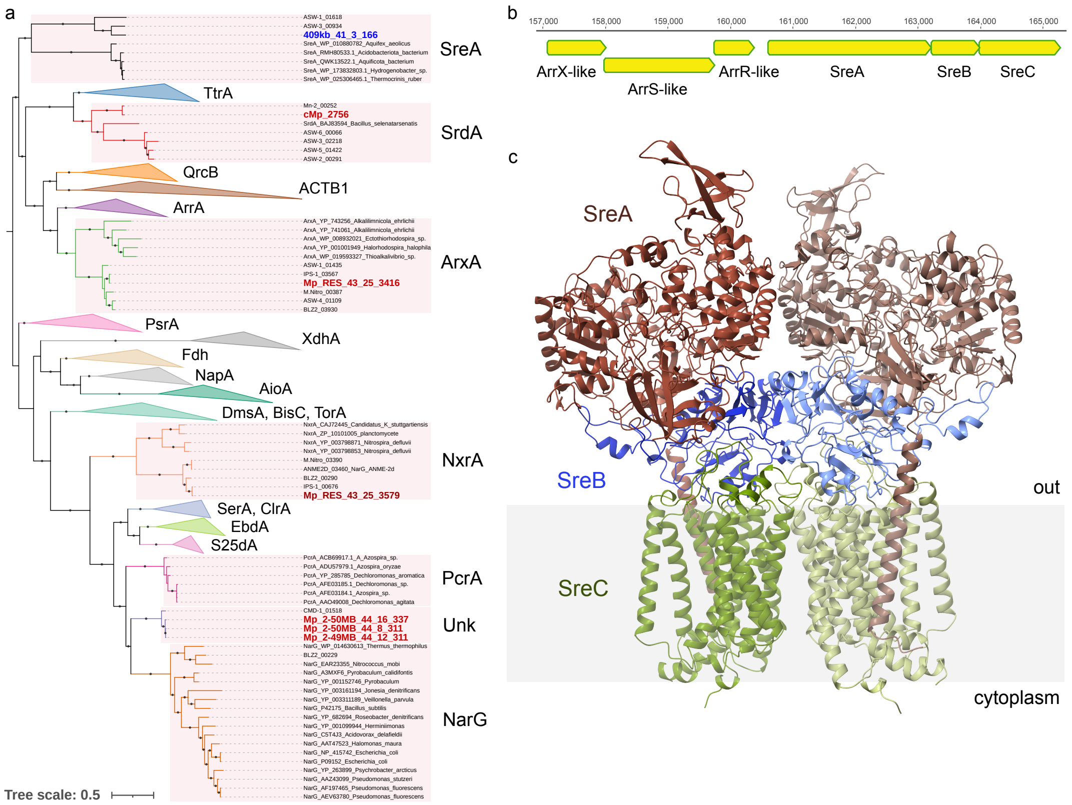


**Supplementary Figure 13. DMSO family protein encoded in jumbo ECEs.** **(a)** Phylogeny of the catalytic subunits. The sequence in blue is found in the jumbo ECE and those in red are in *Methanoperedens* recovered in the study. Bootstrap values were calculated based on 1000 replicates and labeled as greater than 80%. SreA, sulfur reductase; TtrA tetrathionate reductase; SrdA, selenate reductase; QrcB, quinone reductase; ACTB1, alternate complex III subunit B; ArrA, arsenate reductase; ArxA, arsenite oxidase; PsrA, polysulfide reductase; XdhA, xanthine dehydrogenase; Fdh, formate dehydrogenase; NapA, periplasmic nitrate reductase; AioA, arsenite oxidase; BisC, biotin sulfoxide reductase; DmsA, DMSO reductase; TorA, trimethylamine N-oxide reductase; NxrA, nitrite oxidoreductase; SerA, selenate reductase; ClrA, chlorate reductase; EbdA, ethylbenzene dehydrogenase; S25dA, C25 dehydrogenase; PcrA, perchlorate reductase; NarG, respiratory nitrate reductase; Unk, unknown function. **(b)** Gene cluster of the ECE sulfur reductase. **(c)** Predicted heterodimeric structure of the ECE SreABC complex colored by chains. The ipTM and pTM values are 0.82 and 0.83.

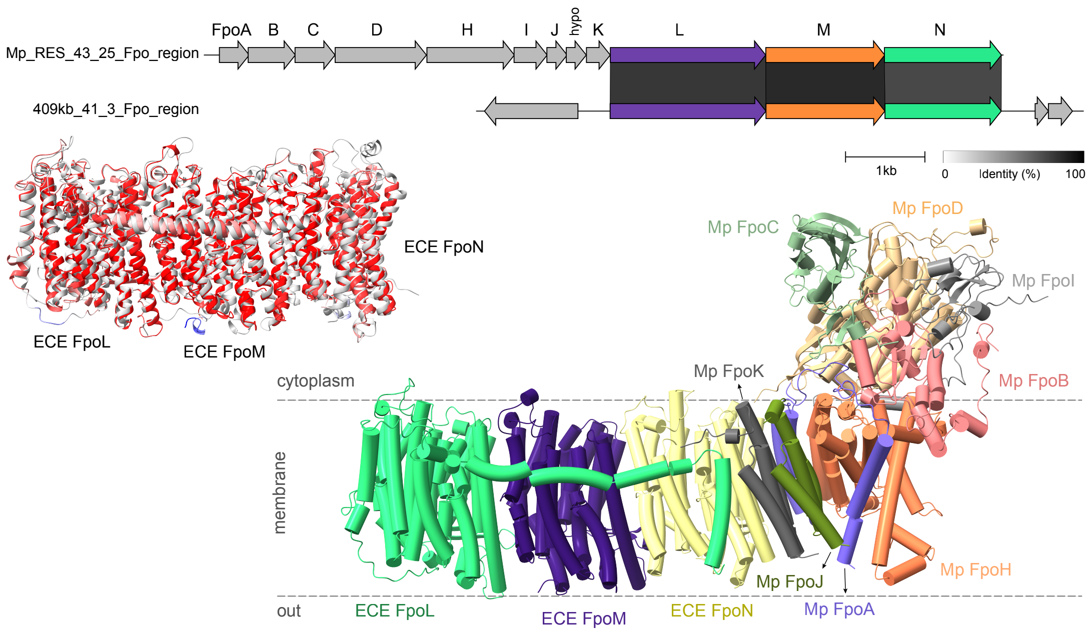


**Supplementary Figure 14. Structural analysis of the Fpo complex encoded by a jumbo ECE.** Homologous genes from the 409kb_41_3 ECE and *Methanoperedens* are connected based on pairwise identity > 30%. Structures of the ECE FpoL, M, and N were predicted individually, colored by pLDDT, and superimposed to the closest reference (8E9I in gray). Then the ECE FpoLMN were co-folded with the *Methanoperedens* FpoABCDHIJK and colored by chains. The ipTM and pTM values of the complex prediction are 0.79 and 0.80.

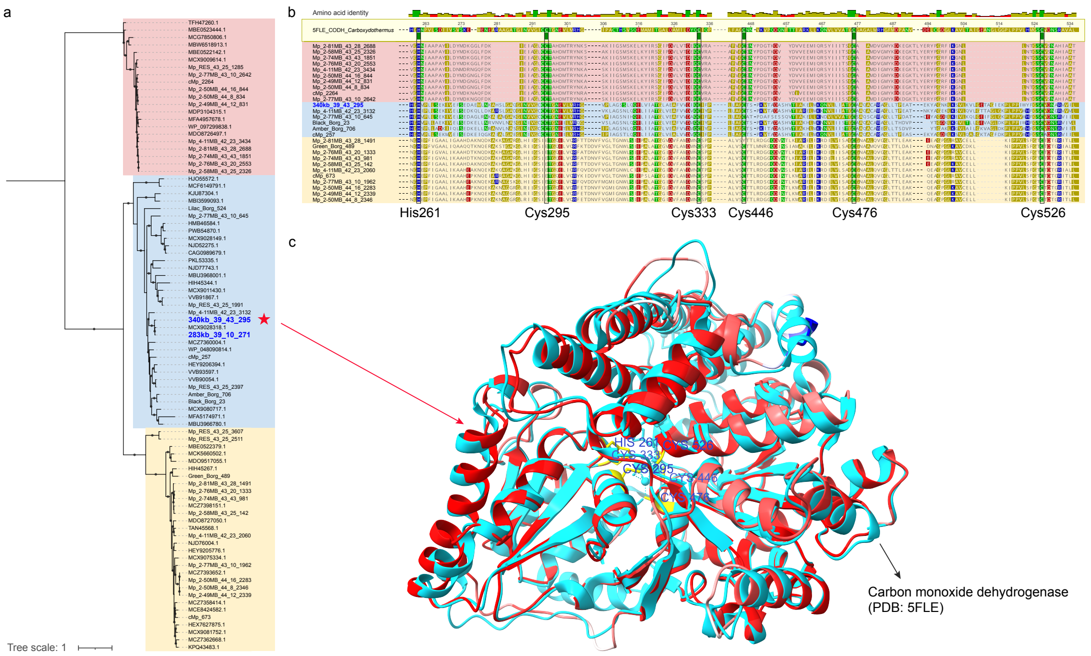


**Supplementary Figure 15. Sequence and structural analyses of carbon monoxide dehydrogenase (CODH). (a)** Phylogeny of CODH in *Methanoperedens* and jumbo ECEs. The ECE sequences are labeled by stars. Dots on branches indicate the same topology observed in > 80% of 1000 re-samplings. **(b)** Alignment of putative CODH sequences. Colored shadows correspond to sequence locations in **(a).** Six residues that coordinate the Ni-Fe-S cluster at the active site are highlighted by boxes. **(c)** Structure prediction and superimposition of CODH from the ECE 340kb_39_43. The query structure is colored based on the pLDDT score. Ni-Fe-S cluster-binding sites are labeled yellow in the experimentally resolved reference.

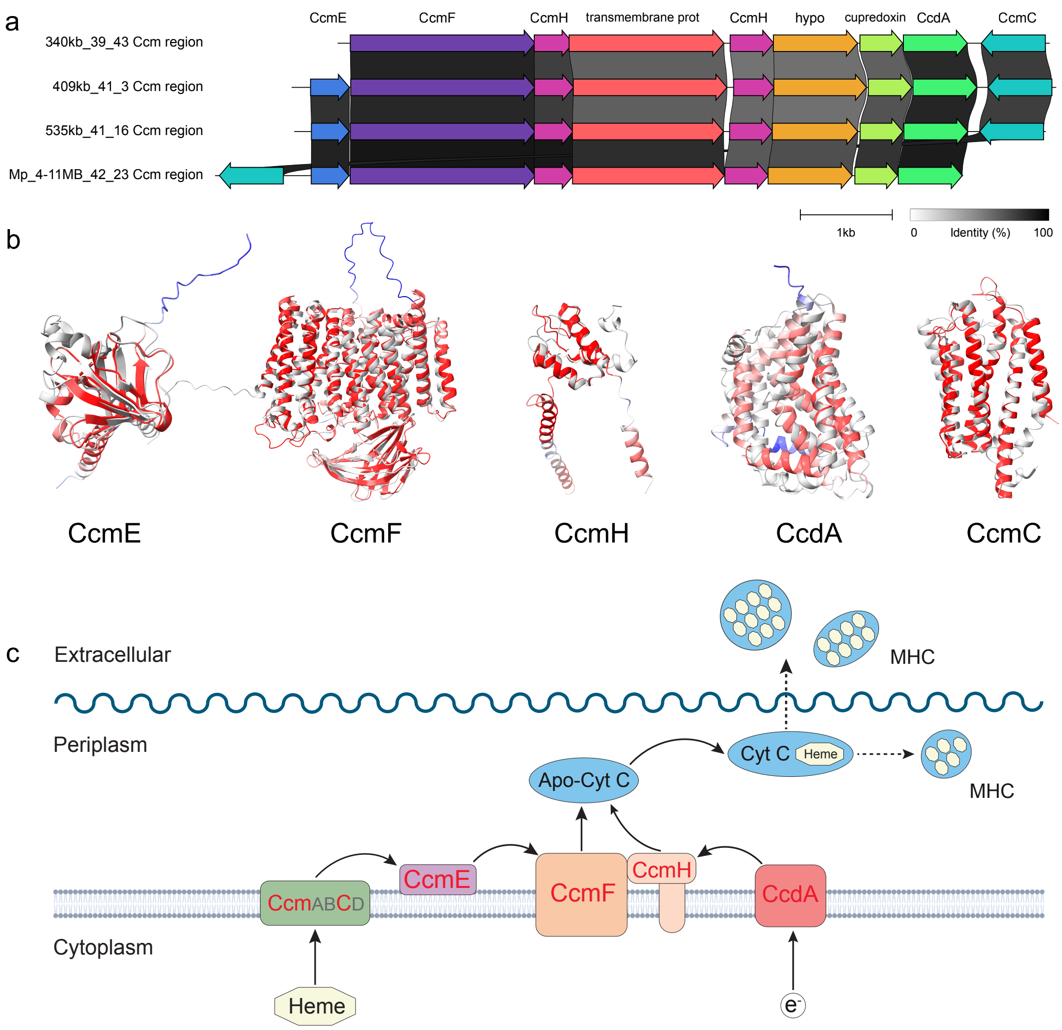


**Supplementary Figure 16. Proteins responsible for cytochrome *c* maturation.** (**a**) Sequence comparison of genes encoding the cytochrome *c* maturation (Ccm) system I. CcmE is not encoded in the jumbo ECE 340kb_39_43. Note CcmC is not adjacent to CcdA or CcmE in genomes; they were placed together for visualization purposes. (b) Structural analysis of Ccm-related proteins in 535kb_41_16. Predicted structures are colored based on the pLDDT score and superimposed with the corresponding best matching references (colored in gray). The references are *E. coli* CcmE (PDB: 8CE8_E), *Thermus thermophilus* CcmF (PDB: 6ZMQ), *Pseudomonas aeruginosa* CcmH (PDB: 2HL7), *Bacillus halodurans* CcdA (Swiss-Prot: Q9KDL8), and *E. coli* CcmC (PDB: 7F02_C). (**c**) Complex pathway for cytochrome *c* maturation. ECE proteins are highlighted in red.

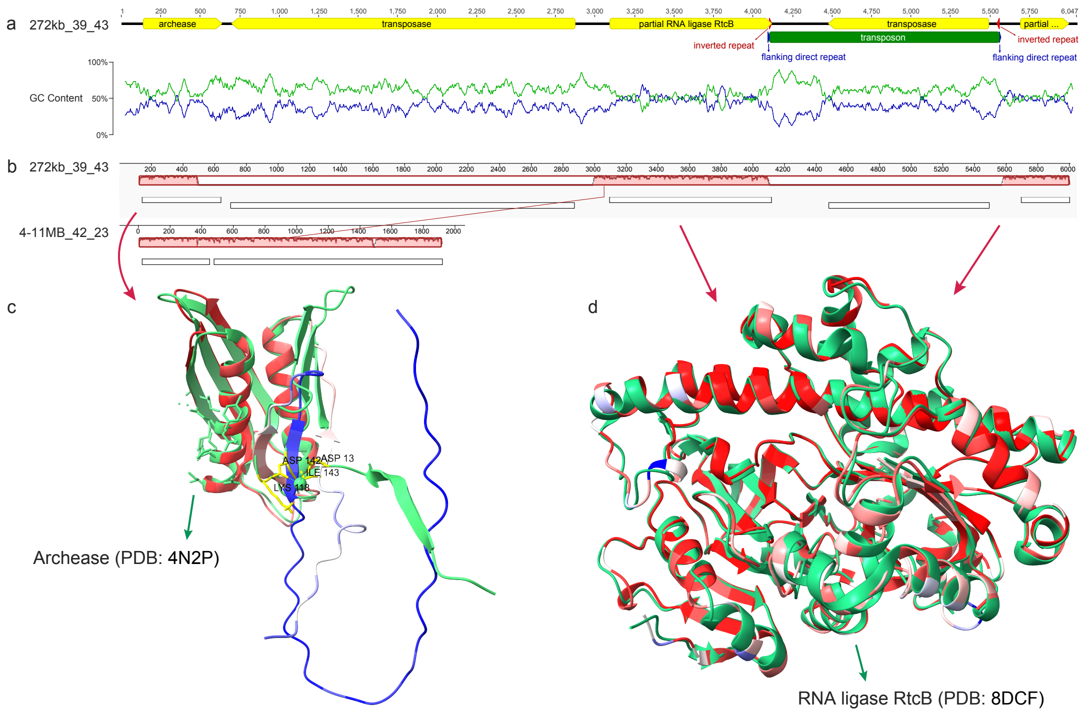


**Supplementary Figure 17. Coordinated proteins for RNA ligation. (a)** Genomic context of archease and tRNA ligase (RtcB). Yellow arrows indicate predicted genes. The green bar indicates the putative transposon terminated by inverted repeats (red arrows) and flanked by short direct repeats (purple arrows). **(b)** Mauve alignment of genomic regions from the jumbo ECE 272kb_39_43 and the *Methanoperedens* 4-11MB_42_23. **(c-d)** Structure prediction and superimposition of archease and reconstructed tRNA ligase from the 272kb_39_43. Structures are colored based on the pLDDT score. Residues of Ca-binding sites in the reference archease and our query protein are ASP13 -> ASP13, LYS118 -> LYS117, ASP142 -> GLN130, and ILE143 -> ILE131 **(c)**. Both terminals of the query protein have low confidence in structure prediction. **(d)** The entire transposon inserted within the ECE tRNA ligase gene was identified by locating the inverted repeats inside and the direct repeats flanking the transposon. The intact tRNA ligase is reconstructed by removing the transposon and one copy of the direct repeat caused by homologous DNA repair during transposition.

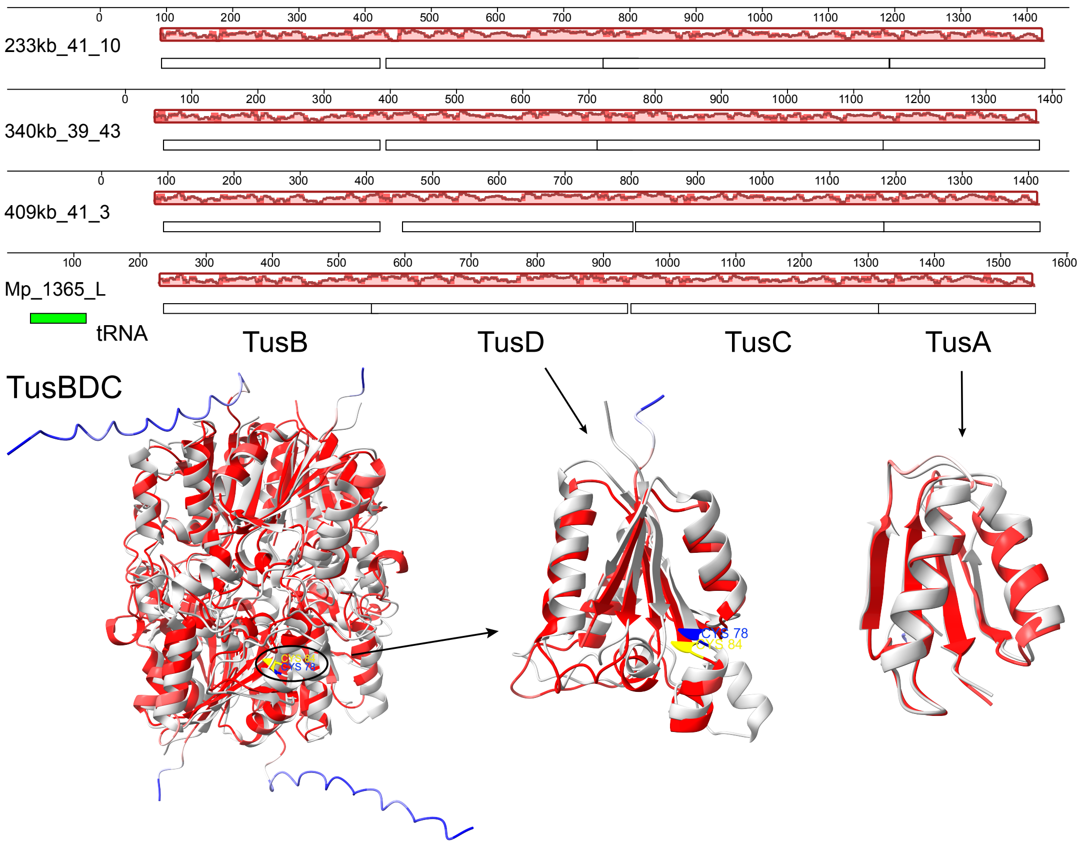


**Supplementary Figure 18. Proteins for sulfur relay in tRNA thiolation.** Mauve alignment was performed on the genomic regions from three jumbo ECEs and one *Methanoperedens*. The representative ECE TusA from 340kb_39_43 is structurally folded alone, and TusBDC are co-folded in a heterohexamer by AlphaFold3. The resulting ipTM and pTM scores of the TusBDC complex are 0.93 and 0.92. All the structures are colored based on the pLDDT score and are superimposed with the best matches from the PDB database (gray; 2D1P and 3LVK). The active site in the TusD subunit participating in sulfur transfer is highlighted in blue for the reference protein (Cys78) and in yellow for the ECE protein (Cys84).

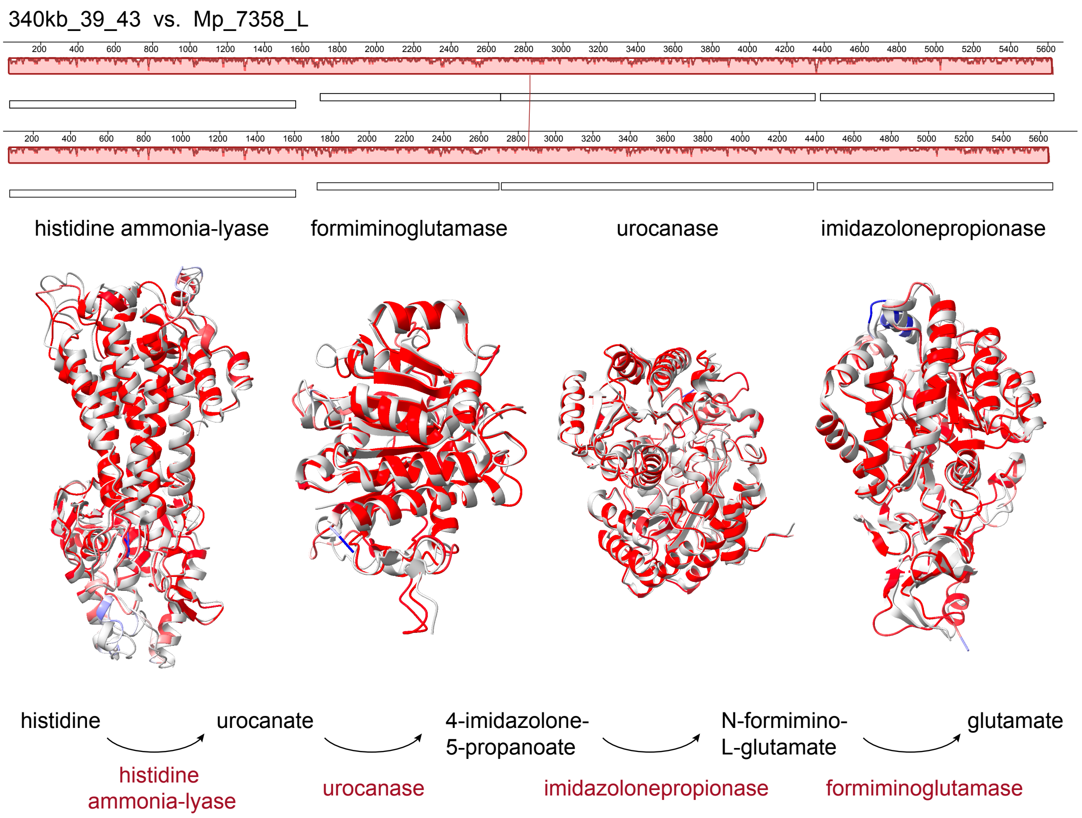


**Supplementary Figure 19. Gene clusters converting histidine to glutamate.** The top one in the Mauve alignment is from the jumbo ECE 340kb_39_43, and the bottom one is from the *Methanoperedens* Mp_7358_L. Structures are predicted for each protein, colored by pLDDT score, and superimposed with the best matches from the PDB database (gray; 1GKJ, 4MXR, 2FKN, and 2G3F, respectively). The reaction from histidine to glutamate is depicted below the structures with the responsible enzymes labeled in red.

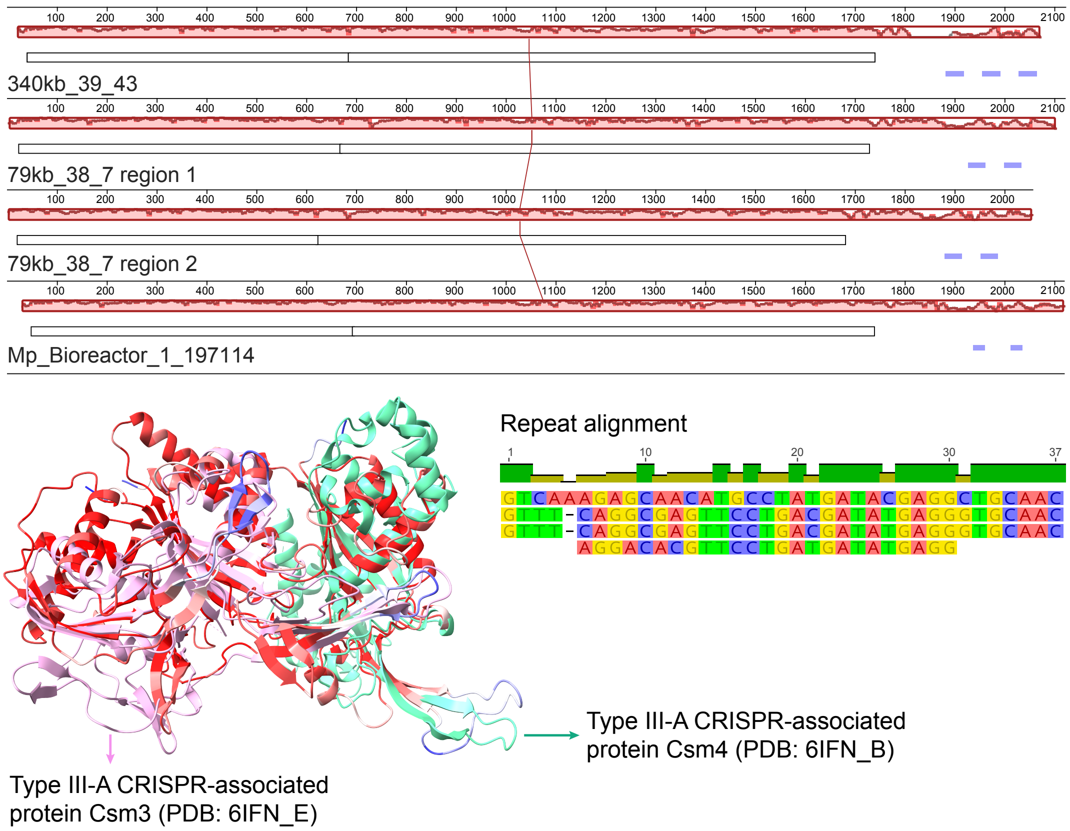


**Supplementary Figure 20. Comparison of Type III-A CRISPR-associated Csm3-Csm4 regions.** The top three genomic regions in the Mauve alignment are from ECEs, and the bottom one is from *Methanoperedens*. The proteins from 340kb_39_43 are co-folded using AlphaFold3, and the resulting ipTM and pTM scores are 0.86 and 0.89. The structures are colored based on the pLDDT score and are superimposed with the best matches from the PDB database (pink and green). Adjacent repeats (blue bars) are aligned following the genome order in the Mauve alignment.

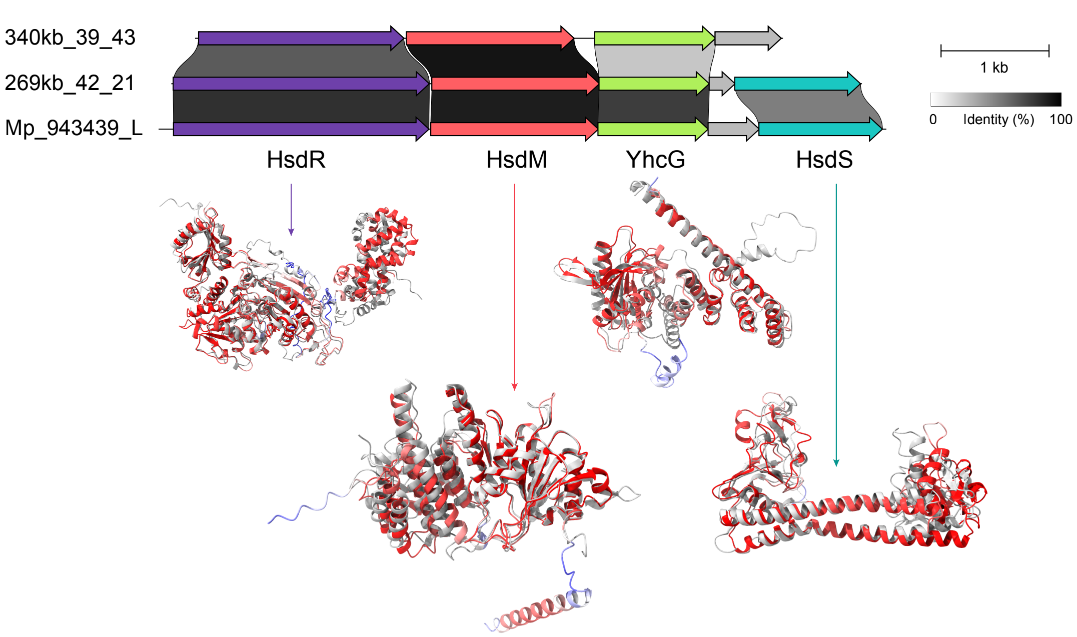


**Supplementary Figure 21. Type I restriction-modification system in the jumbo ECEs and *Methanoperedens*.** Homologous genes are connected based on pairwise identity > 30%. Predicted structures of proteins from 269kb_42_21 are colored by pLDDT scores and superimposed with best matches from the PDB and AFDB-SWISSPROT databases (gray). References for HsdR, HsdM, YhcG, and HsdS are Q07736 (SwissProt), 3UFB (PDB), P45423 (SwissProt), and 5YBB (PDB), respectively.

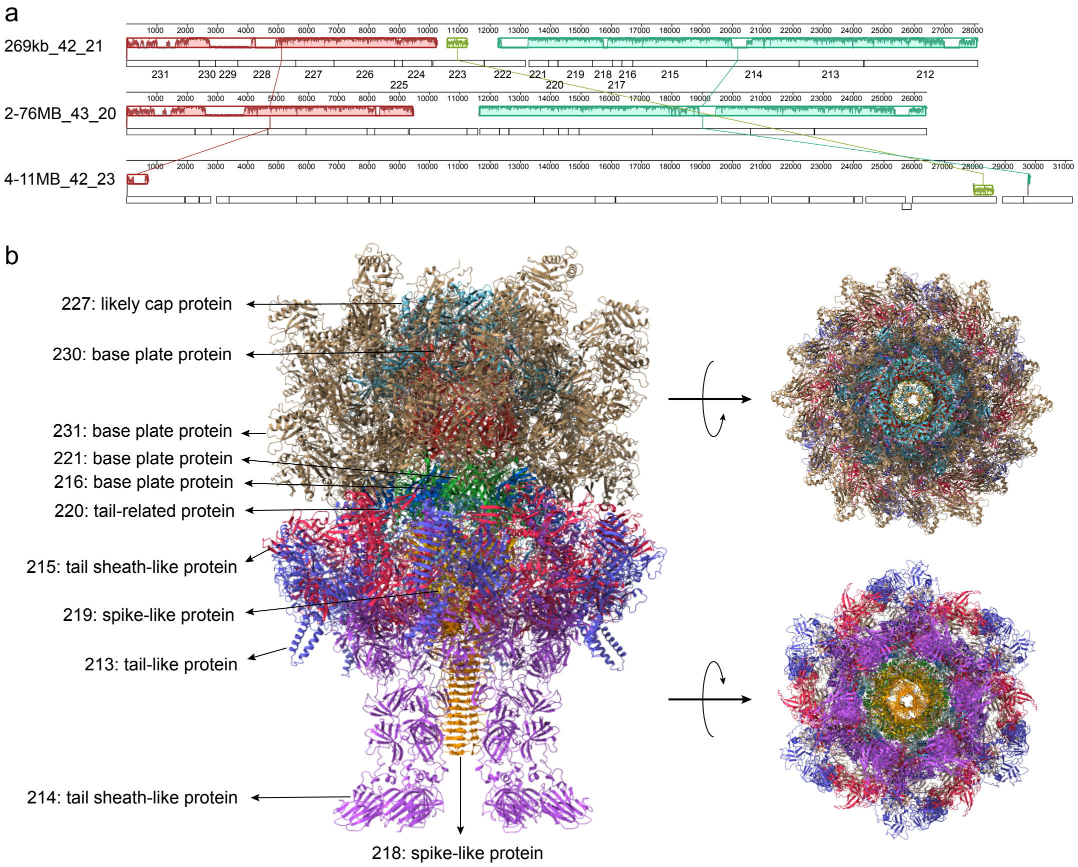


**Supplementary Figure 22. Comparison and structure prediction of extracellular contractile injection systems (eCIS). (a)** Mauve alignment of eCIS regions from *Methanoperedens* and the ECE 269kb_42_21. Note Mp_4-11MB_42_23 is the predicted host by transposon linkage. **(b)** Predicted structure of the 269kb_42_21 by (co-)folding and assembling individual proteins with PDB 7B5H as the template. Structures are colored by chain. Top and bottom views are shown on the right. Proteins that are assembled in the complex are labeled. Some are not assembled, including proteins 212 (tailspike-like protein), 217 (no annotation), 222-224 (eCIS-associated, DUF4157 domain-containing toxins), 225 (no annotation), 226 (ATPase), 228 (no annotation), and 229 (likely baseplate hub assembly protein).

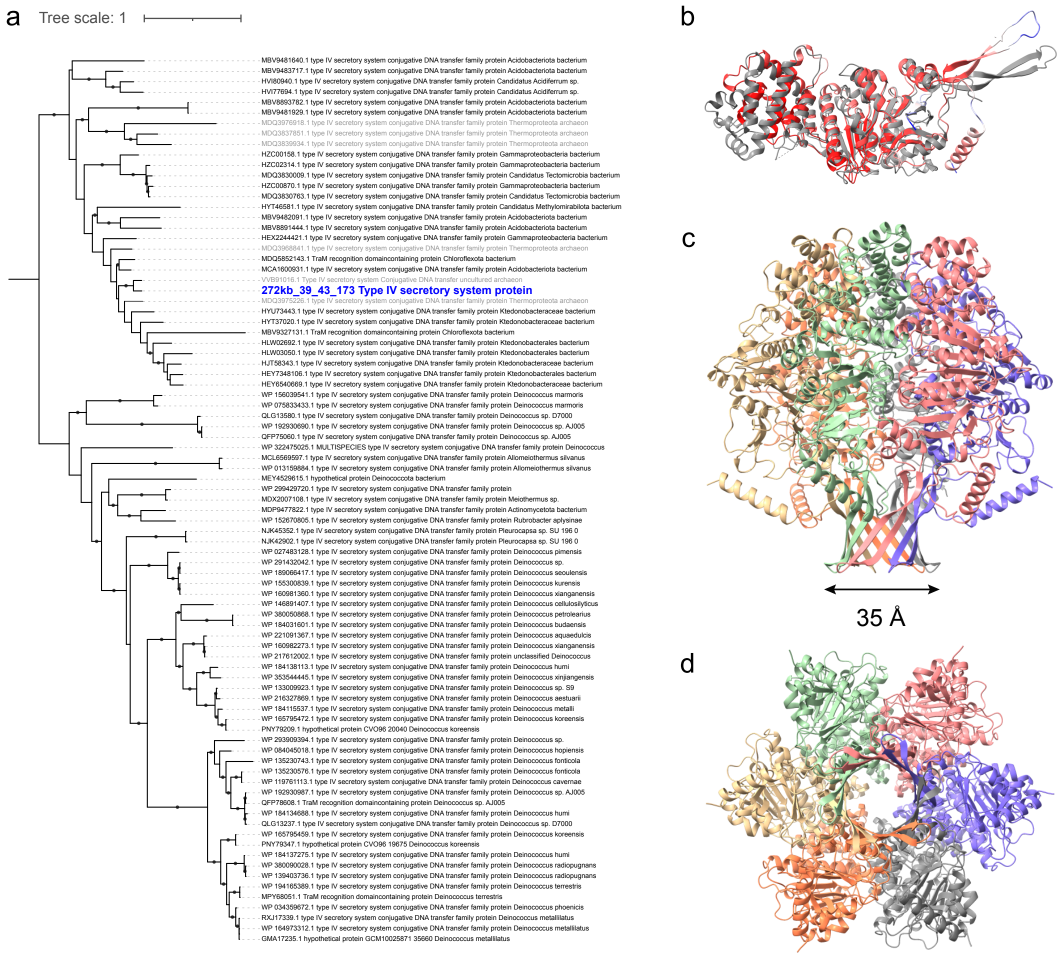


**Supplementary Figure 23. Sequence and structural analyses of Cagβ-like Type IV secretory protein in 272kb_39_43. (a)** Phylogeny of Cagβ-like Type IV secretory proteins. The sequence labeled in bold blue is from the jumbo ECE, and others are recruited from the NCBI database by blasting. Sequences in gray are noted as archaeal, but the corresponding scaffolds that harbor these proteins are too short (< 5 kb) to confidently classify. All other sequences are bacterial. Dots on branches indicate the same topology observed in > 80% of 1000 re-samplings. **(b)** Structure prediction and superimposition of the Cagβ-like Type IV secretory protein from 272kb_39_43. The query structure is colored based on the pLDDT score and the closest matching structure is in gray (PDB: 8DOL). **(c-d)** Side and membrane views of the predicted homohexamer of the ECE Cagβ-like Type IV secretory protein by AlphaFold3. The resulting ipTM and pTM values are 0.73 and 0.78. Structures are colored by chain. The diameter of the central ring channel is measured and labeled using ChimeraX.

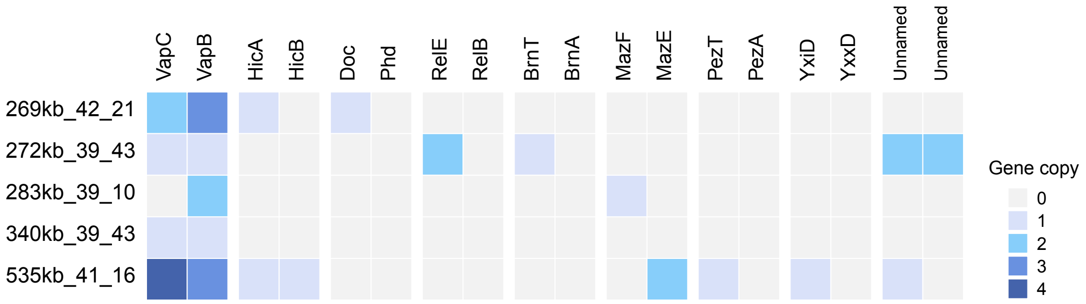


**Supplementary Figure 24. Toxin-antitoxin (TA) systems in the jumbo ECEs.** Colors indicate the gene copies in each genome.
